# High photosynthesis rate in the selected wild rice is driven by leaf anatomy mediating high Rubisco activity and electron transport rate

**DOI:** 10.1101/754887

**Authors:** Jyotirmaya Mathan, Anuradha Singh, Vikram Jathar, Aashish Ranjan

## Abstract

The importance of increasing photosynthetic efficiency for sustainable crop yield increases to feed the growing world population is well recognized. The natural genetic variation for leaf photosynthesis in crop plants is largely unexploited for increasing genetic yield potential. The genus *Oryza*, including cultivated rice and wild relatives, offers tremendous genetic variability to explore photosynthetic differences, and underlying biochemical, photochemical, and developmental bases. We quantified leaf photosynthesis and related physiological parameters for six cultivated and three wild rice genotypes, and identified photosynthetically efficient wild rice accessions. Fitting *A*/*C_i_* curves and biochemical analyses showed that the leaf photosynthesis in cultivated rice varieties, IR64 and Nipponbare, was limited due to leaf nitrogen content, Rubisco activity, and electron transport rate compared to photosynthetically efficient accessions of wild rice *Oryza australiensis* and *Oryza latifolia*. The selected wild rice accessions with high leaf photosynthesis per unit area had striking anatomical features, such as larger mesophyll cells with more chloroplasts, fewer mesophyll cells between two consecutive veins, and higher mesophyll cell and chloroplast surface area exposed to intercellular space. Our results show the existence of desirable variations in Rubisco activity, electron transport rate, and leaf anatomical features in the rice system itself that could be targeted for increasing the photosynthetic efficiency of cultivated rice varieties.

**Highlight:** Distinct leaf biochemical, photochemical, and developmental features contribute to efficient photosynthesis in the selected wild rice accessions that could potentially be exploited to increase rice leaf photosynthesis.

## Introduction

Photosynthesis, the key biochemical process fixing atmospheric CO_2_ in the form of carbohydrates, is a multifaceted process that has contributions from leaf physiological, biochemical, and developmental features (Long *et al*., 2006; Mathan *et al*., 2016). Increasing photosynthetic efficiency of crop plants is suggested to be an important strategy for sustainable increases in crop yield potential and productivity (Zhu *et al*., 2010; Lawson *et al*., 2012; Ort *et al*., 2015; Simkin *et al*., 2019). The extensive biochemical understanding of photosynthesis over the years has highlighted the potential biochemical targets, such as decreasing Rubisco oxygenase activity, increasing RuBP regeneration, and suppressing photorespiration, to improve photosynthetic efficiency (Horton, 2000; Long *et al*., 2006; Murchie and Niyogi, 2011). *in vivo* Rubisco activity (*V_cmax_*), maximum electron transport rate (*J_max_*) determining the regeneration of ribulose phosphate (RuBP) for Rubisco, and the maximum rate of triose phosphate utilization (*TPU*) as the three major biochemical limitations to leaf photosynthesis (Farquhar *et al*., 1980; Sharkey, 1985; von Caemmerer, 2000). The response of net CO_2_ assimilation under saturating light to intercellular CO_2_ concentrations (*A*/*C_i_* curve) is extensively used to deduce these biochemical limitations to efficient leaf photosynthesis (Long and Bernacchi, 2003; Sharkey *et al*., 2007). However, the limited characterization of genetic variability of these biochemical limitations has restricted the potential of increasing leaf photosynthesis in crop improvement.

Leaf photosynthesis limitation due to Rubisco activity, which primarily occurs at low intercellular CO_2_ concentration (< 200 µmol mol^-1^), is due to the low activity of Rubisco along with competing carboxylation and oxygenation reactions mediated by the enzyme (von Caemmerer, 2000; Parry *et al*., 2007; Sharkey *et al*., 2007). Exploration of natural variation in Rubisco activity, as well as developmental and metabolic features mediating increased concentration of CO_2_ and RuBP at the active site of Rubisco, are possible strategies to overcome this limitation of photosynthesis (Parry *et al*., 2007; Parry *et al*., 2013; Lin *et al*., 2014; Sharwood, 2017). Organization of LHCs, the quantum efficiency of the photosystem, and photochemical and non-photochemical quenching are key factors for determining photosynthetic limitation due to electron transport rate and RuBP regeneration that operates at CO_2_ concentration > 300 µmol mol^-1^ (Murchie and Lawson, 2013; Wu *et al*., 2019). *TPU* drives sucrose synthesis from primary photosynthates, and *TPU*-limited photosynthesis primarily operates at high CO_2_ concentrations that often determine *A_max_* (Sharkey *et al*., 2007; Fabre *et al*., 2019). The genotypic variations for these limitations in crop plants have been attributed to photosynthetic differences (Acevedo-Siaca *et al*., 2020; De Souza *et al.,* 2020). For example, Acevedo-Siaca *et al*. (2020) recently investigated the variation among rice accessions to show *V_cmax_* as the primary limitation to photosynthetic induction. Therefore, understanding these biochemical limitations in cultivated crop varieties will provide the potential ways to improve photosynthetic efficiency.

In addition to the biochemical limitations, stomatal conductance (*g_s_*) and mesophyll conductance (*g_m_*) that affect leaf gaseous exchange are critical determinants of leaf photosynthesis (Farquhar and Sharkey, 1982; Adachi *et al*., 2013; Gago *et al*., 2016). Leaf developmental features, besides affecting the leaf gaseous exchange, influence light harvesting and distribution for photosynthesis (Terashima *et al*., 2011; Mathan *et al*., 2016). Flag leaf width (FLW), flag leaf thickness (FLT), and specific leaf area showed a positive correlation with crop photosynthetic efficiency (Liu *et al*., 2015). Anatomical features such as size, lobbing, and surface area of mesophyll cells are critical for proper light distribution and CO_2_ diffusion within a leaf (Flexas *et al*., 2008; Sage and Sage, 2009; Tholen *et al*., 2012). Furthermore, the surface area of mesophyll cell (*S_mes_*) and chloroplast (*S_c_*) exposed to intercellular spaces are the key determinants of *g_m_* (Scafaro *et al*., 2011). Similarly, vein density and interveinal distance influence CO_2_ fixation efficiency (Amiard *et al*., 2005; Brodribb *et al*., 2007). Thus, identifying the exploitable desirable variations in leaf morphological and anatomical features could help in optimizing crop photosynthetic efficiency (Evans, 2013).

Rice, with two cultivated species: *Oryza sativa* (Asian rice with two subspecies: *indica* and *japonica*) and *Oryza glaberrima* (African rice), and many wild rice species, provides an excellent system to investigate the genetic diversity for crop improvement. Rice domestication, which primarily targeted grain features and plant architectural traits, significantly reduced the genetic diversity for many favourable traits in cultivated rice varieties (Huang *et al*., 2012; Chen *et al*., 2021). Wild rice species may harbour more natural allelic diversity than cultivated varieties, and hence mining hidden useful variants from wild species could be an important strategy in rice improvement. Wild rice species are already shown to be reservoirs of many useful traits, particularly for biotic and abiotic stress resistance, which have been used in rice breeding programs (Sanchez *et al.,* 2013; Atwell *et al*., 2014; Wing *et al*., 2018). Interestingly, some of the wild rice accessions of *O. rufipogon*, *O. latifolia*, and *O. australiensis* are reported to accumulate higher biomass than cultivated varieties, and introgression from a wild *Oryza* species to cultivated rice has also been shown to increase biomass (Yeo *et al*., 1994; Zhao *et al*., 2008). Moreover, a few wild rice accessions are also reported to have high leaf photosynthesis rate than the cultivated varieties (Zhao *et al*., 2010; Kiran *et al*., 2013). Thus, wild relatives of rice could be a valuable resource for increasing leaf photosynthetic efficiency of cultivated varieties. However, a systematic effort to identify photosynthetically efficient wild rice accessions along with biochemical, photochemical, and developmental traits promoting high photosynthesis rate in those accessions is warranted.

We examined a set of cultivated and wild rice accessions to find out photosynthetically efficient wild accessions. We identified that the Rubisco activity and electron transport rate limit leaf photosynthesis in the cultivated varieties compared to the selected wild accessions. Photochemical and developmental differences, further, corroborated the leaf photosynthesis differences across the selected cultivated and wild rice.

## Materials and methods

### Plant material and growth conditions

Nine rice genotypes, consisting of four varieties of *Oryza sativa* ssp. *indica* (IR 64, Swarna, Nagina 22, and Vandana), a variety of *Oryza sativa* ssp. *japonica* (Nipponbare), African cultivated rice *Oryza glaberrima*, and three wild rice species, *Oryza rufipogon*, *Oryza latifolia*, and *Oryza australiensis* were used for the study (Supplementary Table S1). Seeds were germinated on germination paper for seven days at room temperature and then transferred to Hoagland solution in test tubes for two weeks of hydroponic growth. Next, fifteen to twenty individual plants of each genotype were transplanted in the research field of the National Institute of Plant Genome Research, New Delhi (average air temperature > 25 °C, 70-80% humidity, and more than 100 cm annual rainfall). In addition, five plants of each genotype were transplanted in the plastic pots, one plant per pot (15 cm diameter × 14 cm height), and were grown in randomized complete block design under controlled growth chambers with re-randomization every seventh day to control the confounder. Plants were supplied with the same amount of Hoagland solution weekly. The growth conditions were maintained as 28/22°C day and night air temperature, 70-80% relative humidity, a photoperiodic cycle of 14h:10h (light:dark), and 1000 µmol m^-2^s^-1^ light intensity. The transition from dark to day light and temperature conditions was incremental. The supply of nutrients, including nitrogen, was controlled for each genotype in both field and controlled growth chamber experiments.

### Photosynthesis rate and related physiological trait measurements

Both field-grown plants, as well as chamber-grown plants, were used for detailed quantification of photosynthesis rate and related physiological traits. Leaf photosynthesis of field-grown plants was measured at ten days after heading from the middle widest portion of fully expanded flag leaves using LI-6400XT portable photosynthesis system using 2 × 3 cm standard leaf chamber (LI-6400, LI-COR, Inc). For the accessions with leaf width less than 2 cm, the leaf chamber was filled using two flag leaves carefully without any overlap (Busch, 2018). For all measurements, the leaf chamber was maintained at ambient atmospheric CO_2_ concentration (*C_a_*, 400 μmol mol^-1^), natural light irradiance (1400-1500 µmol m^-2^s^-1^), constant airflow (300 µmol s^-1^), 70-80% relative humidity, leaf and air temperature (28-30°C), and leaf-to-air maximum vapor pressure deficit in the range of 1.0-1.5 kPa. Gas exchange measurements were performed between 9.00 h to 11.00 h on clear sunny days. Fifteen flag leaves from different plants were used to quantify net photosynthesis rate (*A*), intercellular CO_2_ concentration (*C_i_*), stomatal conductance to water (*g_s_*), and carboxylation efficiency (*CE*, the ratio of *A* to *C_i_*). Values were recorded when *A*, *g_s,_* and *C_i_* became stable.

*A*, *g_s_*, *C_i_* and *CE* were also quantified from the middle widest part of fully-grown eighth leaves of chamber-grown plants using a 2 cm^2^ leaf chamber with a LED light source (LI-6400-02B; LI-COR, Inc). Leaf chamber was maintained at 1500 µmol m^-2^s^-1^ light intensity, 400 μmol mol^-1^ CO_2_ concentration, 300 µmol s constant airflow, 70-80% relative humidity, leaf and air temperature of 28 and 30 °C, respectively, and leaf-to-air maximum vapor pressure deficit in the range of 1.0-1.5 kPa.

### Estimation of leaf chlorophyll content

Total chlorophyll content (*TCC*) was measured in aqueous 80% acetone following the procedure described in Porra *et al*. (1989). Briefly, flag leaves (2.0 × 1.0 cm) of field-grown plants and fully-grown eighth leaves of chamber-grown plants (five biological replicates per genotypes) were harvested and incubated in 5 ml of ice-cold 80% acetone for 48 h, centrifuged at 5,000 × g for 5 min, and absorbance was recorded at 663 nm (Chl a) and 645 nm (Chl b). The pigment concentration (Chl a + b) was calculated using the method described by Arnon, (1949) and expressed in µmol m^-2^.

### Quantification of leaf photosynthesis (*A*) and stomatal conductance (*g_s_*) at different light intensities

Plants grown under controlled growth conditions were used to examine the effect of light intensity on *A* and *g_s_*. The measurements were performed on the same middle widest part of fully-grown eighth leaves of chamber-grown plants (five biological replicates per genotypes) using a 2 cm^2^ standard leaf chamber with a LED light source (LI-6400-02B; LI-COR, Inc). The leaf chamber was maintained at a constant temperature and relative humidity for all measurements as explained above. The CO_2_ concentration (*C_a_*) was set at 400 µmol mol^-1^. Leaves were initially stabilized at the highest irradiance (i.e. 2000 µmol m^-2^ s^-1^), and then PPFD was gradually decreased in a step-wise manner (1800, 1500, 1200, 1000, 700, 500, 300, 100 µmol m^-2^ s^-1^). Data were recorded at each point once *A* and *g_s_* values were stable.

### Quantification of *A*, *g_s_*, and chlorophyll fluorescence at different CO_2_ concentrations

The effect of varying CO_2_ concentration on *A* and *g_s_*, as well as on leaf chlorophyll fluorescence, was quantified simultaneously using plants grown under controlled growth conditions. Values were recorded from the same middle widest part of fully-grown eighth leaves of chamber-grown plants (five biological replicates per genotypes) under 1500 µmol m^-2^ s^-1^ light conditions intensity using 2 cm^2^ standard leaf chamber fluorometer (LI-6400-40 LI-COR, Inc) according to the method described in Li *et al*. (2009) and Fabre *et al*. (2019). Before measurements, leaves were stabilized at ambient CO_2_ concentration (400 µmol mol^-1^) and saturating light intensity for at least 10 minutes. The concentration of CO_2_ in the leaf chamber (*C_a_*) was decreased from 400 µmol mol^-1^ concentration to 300, 200, 150, 100, 80, µmol mol^-1^ CO_2_, and then increased in a step-wise manner to 400, 600, 800, 1000, 1200, and 1400 µmol mol^-1^ CO_2_. The values of *C_i_* at each CO_2_ concentration for the selected accessions are shown in Fig. S4B. The CO_2_ and H_2_O diffusion through the gasket was corrected according to the manufacturer’s instructions (LI-COR 6400XT manual version 6). The leaf and air temperature, relative humidity, airflow, and vapor pressure deficit were maintained as mentioned above.

For photochemical traits, steady-state fluorescence (*F_s_*) was quantified at different CO_2_ concentrations. Light-adapted maximum fluorescence (*F_m_’*) was determined at each CO_2_ concentration by illuminating 2.0 cm^2^ leaf surface inside the chamber with a saturated pulse of light (about 8000 μmol m^−2^ s^−1^ for 0.8 sec), provided by an array of light-emitting diodes (a mixture of 90% red and 10% blue lights at 640 and 460 nm peak wavelength, respectively). Minimal fluorescence level in the light-adapted state (*F_o_’*) was determined after removal of the actinic light, and illuminating the leaf with 1 sec of far-red light (735 nm). Using *F_s_*, *F_m_*’ and *F_o_’*, we calculated (a) the efficiency of PSII (□*PSII)* (b) the coefficient of photochemical quenching (*qP*), and (c) the photosynthetic electron transport rate (*J_PSII_*_)_ in a light-adapted state using following formulae as described in Murchie and Lawson, (2013).

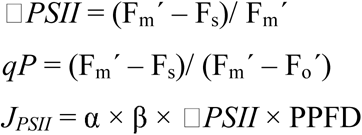

where PPFD is incident photosynthetic photon flux density (i.e., 1500 μ mol m^−2^ s^−1^), β is the photosystem partitioning factor between PSI and PSII for the distribution of light (0.5), and α is the leaf absorbance calculated from total chlorophyll content per unit leaf area (*TCC*, µmol m^-2^) for each rice genotype. *TCC* was quantified from fully grown leaves of growth-chamber grown plants at the same stage when photosynthesis was measured (Supplementary Table S2). α was calculated from *TCC* using the following formula as described in Evans and Poorter, (2001):

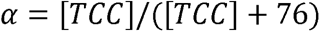

### Quantification of mitochondrial respiration rate in the light (*R*_d_) and the CO_2_ compensation point related to *C_i_* (**Γ*\**)**

R*_d_* and Γ* were quantified at the intersection point of multiple *A*/*C_i_* response curves, conducted at a series of CO_2_ concentrations and multiple light intensities using plants grown under controlled growth conditions (Laisk, 1977; Li *et al*., 2009). *A* was recorded from the same middle widest part of fully-grown eighth leaves of chamber-grown plants using a 2 cm^2^ standard leaf chamber fluorometer (LI-6400-40 LI-COR, Inc). Light intensity inside the leaf chamber was controlled as a series of 100, 200, and 500 μmol m^−2^ s^−1^. At each light intensity, CO_2_ concentration was adjusted in the series of 75, 100, 125, and 150 µmol mol^-1^ inside the chamber. The leaf and air temperature, relative humidity, airflow, and vapor pressure deficit were maintained as explained above. CO_2_ and H_2_O diffusion through the gasket was corrected according to the manufacturer’s instructions (LI-COR 6400XT manual version 6). The value of *A* at the point of intersection of the three *A*/*C_i_* curves represented -*R*_d_, and *C_i_* at the point represented Γ* (Li *et al*., 2009). The representative graphs for the quantification of *R*_d_ and Γ*\** of one plant of each genotype are shown in Supplementary Fig. S5. The average values of *R*_d_ and Γ*\** were obtained from the values of three individual plants of each genotype.

### Quantification of mesophyll conductance (*g_m_*)

Mesophyll conductance (*g_m_*) at ambient CO_2_ concentration (400 µmol mol^-1^) was quantified using the formula *g_m_* = *A*/*C_i_*-*C_c_*), where *C_c_* is the chloroplastic CO_2_ concentration. *C_c_* was calculated according to the variable J method (Harley *et al*., 1992) as:

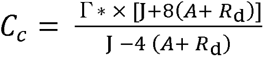

where Γ* is CO_2_ compensation point (μmol mol^-1^), *R*_d_ is mitochondrial respiration rate in the light (μmol m^-2^ s^-1^), and J is the rate of photosynthetic electron transport, which was estimated using chlorophyll fluorescence analysis as *J_PSII_* (described above). Genotype-specific values of variables were used for the calculation of *g_m_* for each genotype.

### Fitting the FvCB model to estimate biochemical limitations

We used the non-linear fitting method explained by Sharkey (2016), version 2 to fit the FvCB model. Three major photosynthetic limitations, the maximum velocity of Rubisco carboxylation (*V_cmax_*), maximum electron transport rate for RuBP regeneration (*J_max_*), and maximum rate of triose phosphate utilization (*TPU*) were estimated simultaneously by minimizing the sum of squares of the residuals. Genotype-specific values of *g_m_*, R*_d,_* and Γ* (as explained above) were used for fitting the model. Rice Rubisco catalytic constants were taken from Perdomo *et al*. (2017). All parameters were scaled to a constant temperature of 25°C. Before fitting the curves, values for each curve were corrected for diffusive leaks between the cuvette and the external environment (Bernacchi *et al*., 2001). Rubisco limitation (*A_c_*), RuBP-regeneration limitation (*A_j_*), and TPU limitation (*A_t_*) to photosynthesis for different accessions were estimated using the following formulae:

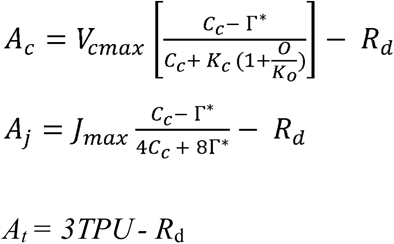

where O is partial pressure for oxygen, and *K_C_* and *K_O_* are Rubisco catalytic constants.

### Rubisco activity assay

Rubisco activity assay was performed following the procedure described in Kubien *et al*. (2011) and Sharwood *et al*. (2016). Leaf tissues (3.6 × 1.0 cm) were excised from the same middle widest part of the same leaves of field-grown plants, from where photosynthesis data were recorded, and homogenized in extraction buffer [EB, 5 mM DTT, 10 mg ml^-1^ PVPP, 2 mg ml^-1^ BSA, 2 mg ml^-1^ PEG, 2% (v/v) Tween-80, 12 mM 6-aminocaproic acid, 2.4 mM benzamidine in HEPES stock) at 4 °C. The homogenate was centrifuged for 1 min at ∼16,000 × *g* and the pellet was discarded. To activate rubisco enzyme, leaf extracts were incubated with BMEC (100 mM Bicine-NaOH, pH 8.2, 20 mM MgCl_2_, 1 mM Na_2_-EDTA and 10 mM NaHCO_3_) in a 9:1 ratio (e.g. 900 μL leaf extract to 100 μL BMEC) at 25 °C for 25 min. To assess the activity of Rubisco from leaf extract, 50 µl of leaf extract was incubated with 1,550 ml of the EME buffer, 40 µl 10 mM of NADH (final concentration 200 µM), 80 µl of 250 mM NaH^12^CO_3_ (10 mM), 200 µl of ATP/phosphocreatine (1 and 5 mM, respectively), 40 µl of coupling enzymes (Carbonic anhydrase 500U, Glyceraldehyde-3-P dehydrogenase 50U, 3-PGA kinase 50U, Triose-P Isomerase/Glycerol-3-P dehydrogenase 400/40U), 40 µl of 25 mM RuBP (500 µM), mixed gently, and decrease of absorbance were monitored at 340 nm continuously for 0 to 1800 sec using a UV–Vis spectrophotometer.

The Rubisco carboxylation rate was determined using the equation:

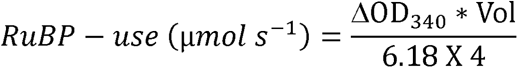

assuming an extinction coefficient of 6.18 µmol^-1^ ml^-1^ cm for NADH oxidation at 340 nm. ΔOD_340_ is the rate of change of absorbance at 340 nm per second, and Vol is assay volume (1 ml). The equation accounts for the four NADH molecules oxidized per RuBP carboxylated by Rubisco.

The activity of Rubisco (*V_cmax_*) is calculated as:

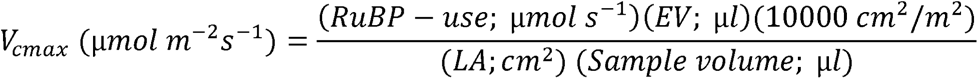

where EV is extraction volume and LA is leaf area. All the chemicals used for the Rubisco activity assay, including RuBP, were commercially procured from Sigma Aldrich (Kubien *et al*., 2011; Sharwood *et al*., 2016).

### Quantification of leaf nitrogen content

Nitrogen content in leaf tissues was calculated by excising 2.0 × 1.0 cm leaf area from the same portion of the leaves where Rubisco activity was performed. The fresh weight of the leaf tissues was determined followed by drying of the tissues for three days at 60 °C. The dried tissues were weighed and ground to a fine powder. The leaf mass per area (LMA) was calculated by dividing the leaf dry weight by leaf area. The nitrogen content of leaves per unit dry weight (*N_mass_*) was then quantified using a Kjeldahl auto-analyzer (Technicon, Tarrytown, New York) as described in Traw and Ackerly (1995). The leaf nitrogen content per unit area (*N_area_*) was derived from N_mass_ and LMA (Pinto *et al*., 2016).

### RNA isolation and expression analysis

Total RNA was isolated from flag leaves in three biological replicates for each genotype, and each biological replicate consisted of tissues from multiple individual plants. Tissues were harvested and immediately frozen in liquid nitrogen. Total RNA extraction (TRIzol reagent), DNase treatment, and cDNA synthesis (RevertAid First-strand cDNA synthesis kit) were performed according to the manufacturer protocol (Thermo Fisher Scientific). The PowerUp™ SYBR® Green fluorescence dye was used for quantitative gene expression analysis on CFX connect Real-time PCR detection system (Bio-Rad). The details of genes used for expression analysis and the primers are presented in Supplementary Table S3 and S4. The specificity of amplified gene products across the selected rice genotypes using the listed primers was viewed through dissociation curve analysis. Before quantitative real-time PCR, the efficiency of each primers pair was evaluated using a standard curve method where four cDNA quantities (1, 10, 50, and 100 ng) from one of the cultivated rice (Nipponbare) was used to construct a standard curve, and efficiency was calculated for each genotype using the formula E = (10^[-1/slope]^-1)*100. Expression analysis of each gene was performed in three biological replicates for each genotype and repeated twice for the reproducibility of the results. We normalized the transcript levels of genes independently to two stable internal controls actin (LOC_Os03g50885.1) and ubiquitin (LOC_Os03g03920) with similar results. Relative expressions of the target genes are presented using both actin and ubiquitin as the internal control following the method by Livak and Schmittgen (2001) (Fig. 4, Supplementary Fig. S9).

### Visualization and quantification of leaf developmental traits

The widest middle part of the flag leaves of 10 -15 field-grown plants was used to measure flag leaf width (FLW) and leaf thickness (FLT). For anatomical traits visualization, transverse sections were cut from the mid-portion of flag leaves (one leaf per plant, five biological replicates per genotype), and stained with toluidine blue with some modification of the method as described in Lux *et al*. (2005). Briefly, chlorophyll pigments of the thin sections were cleared using 85 % (w/v) lactic acid saturated with chloral hydrate, heated on a water bath at 70 °C for 1 h, and washed several times with distilled water. To visualize mesophyll cells, sections were incubated in absolute alcohol overnight and stained with 1% toluidine blue in 1% (w/v) disodium tetraborate for 15 sec. Finally, thin sections were mounted with 50 % glycerol and visualized using a bright field microscope (LMI, UK). The quantification of leaf anatomical features related to the minor vein, mesophyll cell and, bundle sheath cells were performed under a light microscope at 10X and 100X magnification (one leaf per plant, four images per leaf, two on each side of the midrib that consisted of about 3-6 complete cells per image, five biological replicates per genotype) using Fiji-Image J software (Wayne Rasband, National Institute of Health) as described in Chatterjee *et al*. (2016) and Ivanova *et al*. (2018). The mesophyll cell number per unit leaf area, mesophyll cell area, and the mesophyll surface area exposed to the intercellular space (*S_mes_)* were calculated from transverse sections under bright field microscopy as described in Scafaro *et al*. (2011). Total 30 to 40 mesophyll cells per genotype were quantified for the calculation of *S_mes_*.

### Chloroplast ultrastructure

Freshly harvested flag leaf segments (1 × 1 mm; one leaf each from three individual plants per genotype) were fixed under buffer (3% glutaraldehyde in 0.1 M cacodylate buffer, pH 7.2) for 2 h, and then treated with 1% osmium tetroxide overnight at 4 °C. The fixed segments were dehydrated in a graded acetone series and embedded in epoxy resin. Ultrathin sections were cut with diamond knives, stained with uranyl acetate and lead citrate double staining procedure as described by Reynolds, (1963). Chloroplasts of the uppermost part of the leaf sections were viewed using a digital camera (BH-2, Olympus) equipped with transmission electron microscopy at 1000X magnification (JEM 1230; JEOL). The transverse sections prepared for electron microscopy were used to calculate chloroplast area, chloroplast number per unit mesophyll cell area, and the surface area of the chloroplast exposed to intercellular space (S*_c_*) as described in Scafaro *et al*. (2011). 15 to 20 mesophyll cells with clear images of chloroplast cells were quantified for the calculation of *S_c_*.

### Statistical analysis

Differences among the genotypes were tested by one-way analysis of variance (ANOVA) (JMP Pro, version 12.0.1; SAS Institute, Cary, NC, USA). Statistically similar genotypes were grouped according to post-hoc Tukey HSD calculation. Significance test for the genotypes at varying light intensities and CO_2_ concentrations, as well as for Rubisco activity differences was performed using one-way Analysis of Variance (ANOVA). Pearson correlation matrix was calculated based on mean values of each trait for each genotype to evaluate the trait-to-trait associations. The threshold for statistical significance was *P* ≤ 0.05. Linear regression analyses between *V_cmax_*/*J_max_*, *V_cmax_*/*TPU*, and *J_max_*/*TPU* were performed using Microsoft Excel.

## Results

### Quantification of leaf photosynthesis and related physiological features

We observed remarkable differences in leaf photosynthesis (*A*) and related physiological features among the selected cultivated and wild rice accessions (Fig. 1). The mean *A* value of the wild rice species, *O. latifolia*, *O. australiensis*, and *O. rufipogon*, and African cultivated rice *O. glaberrima* were found to be significantly higher than that of cultivated rice varieties, such as IR64 and Nipponbare (Fig. 1A). *O. australiensis* and *O. latifolia* showed the highest *A*, 23.5 and 24.2 µmol CO_2_ m^-2^ s^-1^, respectively, which were 29% to 57% higher compared with the different Asian cultivated varieties. The African cultivated rice *O. glaberrima* showed *A* value intermediate between the wild accessions and Asian cultivated varieties. We also observed significant variations for *A* among the cultivated varieties. For instance, one of the high-yielding *indica* variety Swarna and *japonica* variety Nipponbare displayed 19% and 14% higher *A*, respectively, compared with IR64 (Fig. 1A).

**Fig. 1.**
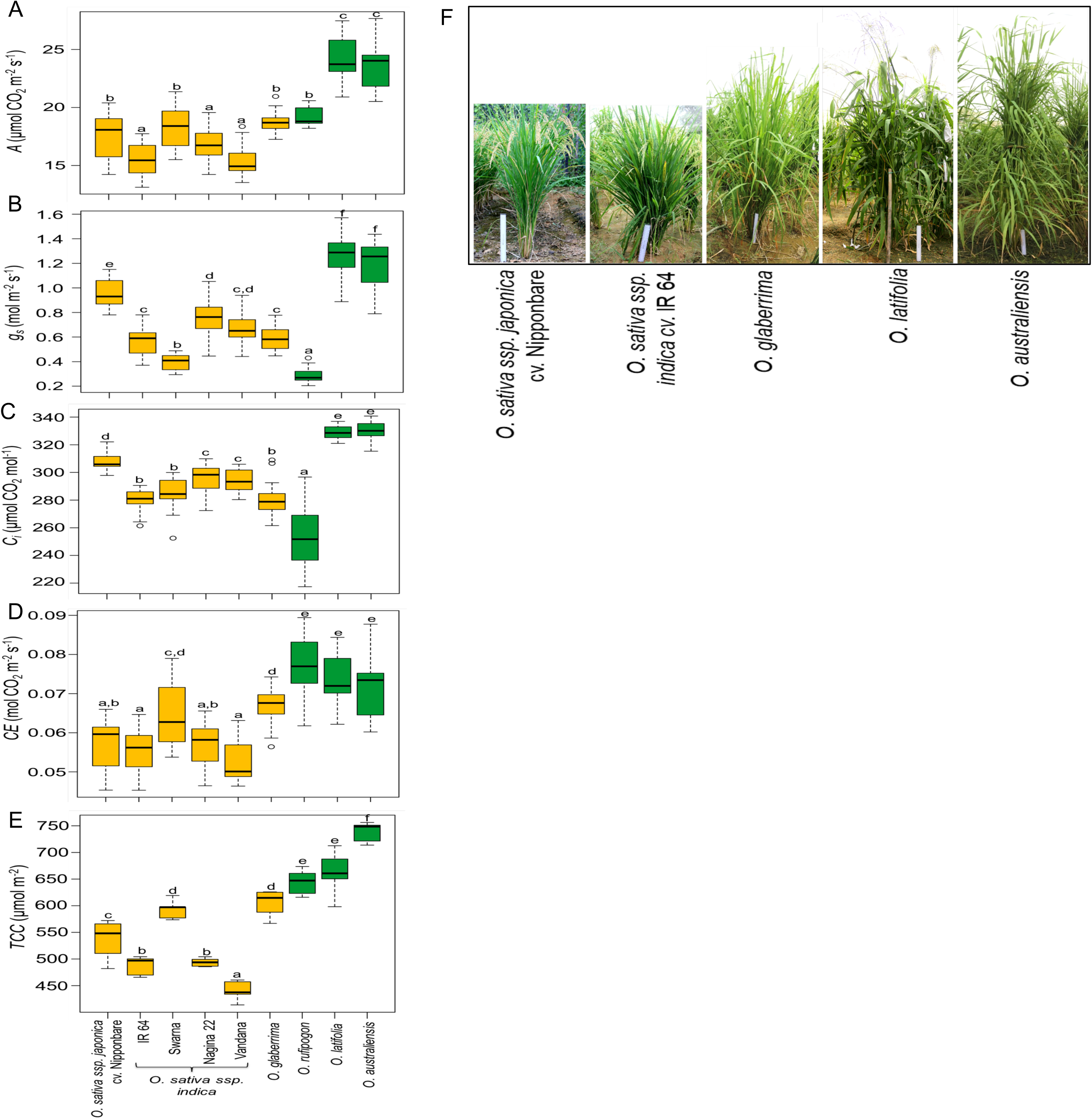
Variations in leaf photosynthesis and related physiological features, as well as morphological differences across the selected cultivated and wild rice accessions. (A-E) Quantification of net photosynthesis per unit area of flag leaf, *A* (A), stomatal conductance to water, *g_s_* (B), intercellular CO_2_ concentration, *C_i_* (C), carboxylation efficiency, *CE* (D), and Total chlorophyll content, *TCC* (E) for six cultivated rice and three wild rice species. Cultivated rice varieties and wild species are represented with yellow- and green-shaded boxes, respectively. Each box plot shows the median and interquartile range with minimum and maximum values of fifteen data points from different plants. Different letters indicate statistical significance according to one-way ANOVA followed by post-hoc Tukey HSD calculation at *P* ≤ *0.05*. (F) Representative images of the selected cultivated and wild rice accessions that captured the entire variation range for the leaf photosynthesis per unit area in the experiment. Shown are the field-grown plants of the selected genotypes. Scale bars represent 30 cm.

We further examined important physiological traits of the selected accessions that are known to influence leaf photosynthesis. As expected, wild rice accessions of *O. latifolia* and *O. australiensis* with the highest *A* showed higher stomatal conductance (*g_s_*), and intercellular CO_2_ concentration (*C_i_*) (Fig. 1B, C). For example, *O. latifolia* and *O. australiensis* accessions showed 1.3 to 3-fold higher *g_s_* compared with different Asian cultivated varieties. In addition, these species along with *O. rufipogon* displayed higher carboxylation efficiency (*CE*) than those of cultivated varieties (Fig. 1D). Among the cultivated varieties, Nipponbare that displayed higher *A* also showed reasonably higher *g_s_* and *Ci* compared with the other cultivated varieties. We also observed significantly higher total chlorophyll content (*TCC*) in flag leaves of the selected wild rice accessions compared with the cultivated varieties. Consistent with the high *A*, wild accessions of *O. latifolia* and *O. australiensis* had 33% and 43% higher *TCC* per unit leaf area, respectively, than IR64. *O. glaberrima*, Swarna, and Nipponbare also had reasonably higher *TCC* than IR 64, Nagina 22, and Vandana (Fig. 1E). Pearson correlation analysis showed a significant positive correlation of *A* with *g_s_* (*r = 0.629*), *C_i_* (*r = 0.570*), *CE* (*r = 0.819*), and *TCC* (*r = 0.912*)*, g_s_* with *C_i_* (*r = 0.965*) and *CE* with *TCC* (*r = 0.906)* at *P* ≤ *0.05* (Supplementary Fig. S1).

We then selected representative cultivated rice accessions (*O. sativa* ssp. *indica* cv. IR 64, *O. sativa* ssp. *japonica* cv. Nipponbare, and African cultivated rice *O. glaberrima*) and wild rice accessions (*O. latifolia* and *O. australiensis)*, which captured the entire variation range for the leaf photosynthesis per unit area in the field experiment, for the quantification of leaf photosynthesis and related physiological traits at the vegetative stage under controlled conditions (Fig. 1F; Supplementary Fig. S2). Similar to our observations at the full heading stage under field conditions, the selected wild rice accessions together with African cultivated rice *O. glaberrima* showed significantly higher *A*, *g_s_*, and *CE* compared to cultivated rice varieties, IR64 and Nipponbare, at the vegetative stage (Table 1). Moreover, we have recently reported consistently higher photosynthesis per unit leaf area in the *O. australiensis* accession compared with Nipponbare at different developmental stages in field conditions (Mathan *et al*. 2021). Taken together, these results showed significant differences in photosynthetic traits across the selected cultivated and wild rice accessions.

**Table 1.** Leaf photosynthesis rate and related physiological traits of the selected cultivated and wild rice accessions under controlled growth conditions. Each value represents mean ± SE, where n = five data points from different plants. Different letters indicate statistical significance according to one-way ANOVA followed by post-hoc Tukey HSD calculation at *P* ≤ *0.05*.

| Genotypes | Cultivated/<br>Wild | $A$<br>($\mu\text{mol CO}_2 \text{ m}^{-2} \text{ s}^{-1}$) | $C_i$<br>($\mu\text{mol CO}_2 \text{ mol}^{-1}$) | $g_s$<br>($\text{mol m}^{-2} \text{ s}^{-1}$) | $CE$<br>($\text{mol CO}_2 \text{ m}^{-2} \text{ s}^{-1}$) |
| --- | --- | --- | --- | --- | --- |
| *O. sativa* cv. Nipponbare | Cultivated | 11.15 ± 1.2a | 309.0 ± 40.5a | 0.22 ± 0.05a | 0.03 ± 0.005a |
| *O. sativa* cv. IR 64 | Cultivated | 11.22 ± 1.1a | 300.9 ± 14.5a | 0.21 ± 0.01a | 0.03 ± 0.008a |
| *O. glaberrima* (IR 102925) | Cultivated | 13.79 ± 0.4b | 282.6 ± 12.9a | 0.28 ± 0.02b | 0.04 ± 0.003a |
| *O. latifolia* (IRGC 99596) | Wild | 18.38 ± 0.1c | 306.2 ± 13.9a | 0.40 ± 0.03c | 0.06 ± 0.001b |
| *O. australiensis* (IRGC 105272) | Wild | 19.18 ± 0.1c | 279.8 ± 21.2a | 0.35 ± 0.02c | 0.06 ± 0.001b |

Next, we examined the effect of different light intensities and CO_2_ concentrations on *A* and *g_s_* of the selected wild and cultivated rice accessions grown under controlled conditions (Fig. S3). *O. latifolia* and *O. australiensis* accessions exhibited higher increases in *A* in response to increasing PPFD and maintained higher *g_s_* at different light intensities compared with the cultivated varieties (Supplementary Fig. S3A and B). *O. latifolia* and *O. australiensis* had significantly higher light-saturated *A_ma_*_x,_ 20.94 and 21.68 µmol CO_2_ m^-2^ s^-1^, respectively than *O. sativa* cv. IR 64 and *O. sativa* cv. Nipponbare (11.89 and 13.02 µmol CO_2_ m^-2^ s^-1^, respectively). Since the differences in the *C_i_* values at different PPFD may confound the photosynthesis rate in different accessions, we quantified the *C_i_* values at each PPFD for the selected accessions. Changes in *C_i_* in response to increasing PPFD during the light response curves were mostly similar across all the genotypes (Fig. S4). The effect of increasing CO_2_ concentration on *A* segregated the selected genotypes under three distinct groups: *O. australiensis* and *O. latifolia* with higher *A*, African cultivated rice *O. glaberrima* with intermediate *A*, and cultivated varieties Nipponbare and IR 64 with lower *A* at different CO_2_ concentration (Fig. S3C). The *g_s_*, however, showed a decreasing trend at increasing CO_2_ concentration, likely due to the closure of stomata (Fig. S3D). Altogether, the selected wild rice accessions of *O. australiensis* and *O. latifolia* were able to maintain higher leaf photosynthesis per unit area at different light intensities and CO_2_ concentrations compared with the selected cultivated varieties.

### Fitting CO_2_ response curves to investigate biochemical limitations *V_cmax_*, *J_max_*, and *TPU*

To investigate the biochemical limitations to photosynthesis across the selected accessions using the FvCB model, we first estimated mesophyll conductance (*g_m_*) at ambient CO_2_ concentration. *g_m_* significantly varied for the selected cultivated and wild rice species (Table 2). *O. australiensis* and *O. latifolia*, with highest *A*, also showed higher *g_m_* than the cultivated varieties, suggesting the key importance of *g_m_* for photosynthetic variations among the genotypes. We also quantified the mitochondrial respiration rate in the light (*R*_d_) and CO_2_ compensation point related to *C_i_* (*Γ\**) for each genotype to fit the FvCB model (Table 2 and Supplementary Fig. S5). *R*_d_ was found to be significantly higher for *O. australiensis* and *O. latifolia* compared to the selected cultivated accessions, whereas no significant difference was observed for *Γ\** across the accessions.

**Table 2.**
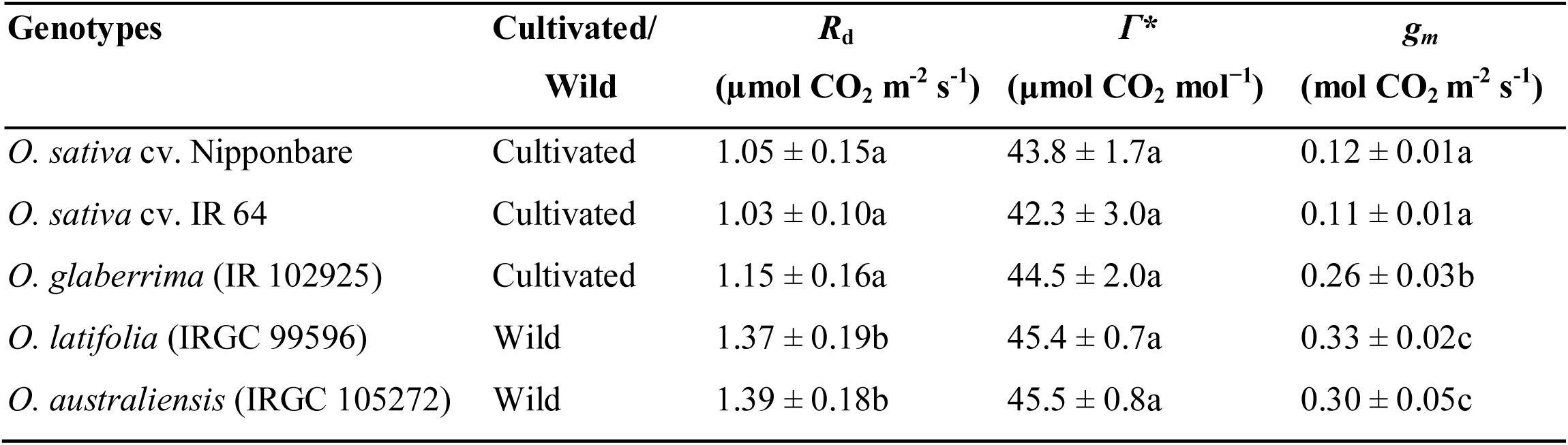
Mitochondrial respiration rate in the light (*R*_d_), CO_2_ compensation point related to *C_i_* (*Γ\**), and mesophyll conductance (*g_m_*) for the selected cultivated and wild rice genotypes. Each value represents mean ± SE, where n = three data points from different plants. Different letters indicate statistical significance according to one-way ANOVA followed by post-hoc Tukey HSD calculation at *P* ≤ *0.05*.

We, then, used carbon dioxide response (*A*/C*_i_*) curves to estimate maxima of Rubisco carboxylation activity (*V_cmax_*), regeneration of ribulose-1,5-biphosphate expressed as electron transport rate (*J_max_*), and triose phosphate utilization (*TPU*) for the selected accessions (Farquhar *et al*., 1980; von Caemmerer, 2000) (Supplementary Fig. S6). CO_2_-saturated *A_max_* for the selected accessions ranged from 21.37 µmol CO_2_ m^-2^ s^-1^ for cultivated variety IR 64 to 36.81 µmol CO_2_ m^-2^ s^-1^ for the wild species *O. australiensis* (Table 3). We found significantly higher values *V_cmax_*, *J_max_*, and *TPU* for the wild species *O. australiensis* and *O. latifolia* compared to cultivated varieties IR64 and Nipponbare, suggesting *V_cmax_*, *J_max_*, and *TPU* together might contribute to differences in *A_max_*. Consistent with the intermediate value of *A_max_* for *O. glaberrima*, the values of biochemical limitations were also intermediate between the wild species and cultivated *indica* and *japonica* varieties. We investigated the relationship among the modelled biochemical limitations to photosynthesis across the selected accessions. A distinct linear relationship was observed between *V_cmax_* and *J_max_* (R^2^ = 0.973), *J_max_* and *TPU* (R^2^ = 0.987), and *V_cmax_* and *TPU* (R^2^ = 0.968) at *P < 0.001*, suggesting the robustness of the modelled limitations (Supplementary Fig. S7). These results, expectedly, indicated the important contribution of biochemical attributes to photosynthetic differences among the selected cultivated and wild rice accessions.

**Table 3.** CO_2_-saturated maximum photosynthesis (*A_max_*), maximum *in vivo* Rubisco activity (*V_cmax_*), maximum electron transport rate (*J_max_*), and maximum rate of triose phosphate utilization (*TPU*) through *A/C_i_* curve analysis. Each value represents mean ± SE, where n = five data points from different plants. Different letters indicate statistical significance according to one-way ANOVA followed by post-hoc Tukey HSD calculation at *P* ≤ *0.05*.

| Genotypes | Cultivated/<br>Wild | $A_{max}$<br>($\mu\text{mol m}^{-2} \text{s}^{-1}$) | $V_{cmax}$<br>($\mu\text{mol m}^{-2} \text{s}^{-1}$) | $J_{max}$<br>($\mu\text{mol m}^{-2} \text{s}^{-1}$) | $TPU$<br>($\mu\text{mol m}^{-2} \text{s}^{-1}$) |
| --- | --- | --- | --- | --- | --- |
| *O. sativa* cv. Nipponbare | Cultivated | 23.99 ± 0.40b | 62.75 ± 0.92b | 124.86 ± 1.37b | 9.10 ± 0.06b |
| *O. sativa* cv. IR 64 | Cultivated | 21.37 ± 0.88a | 51.48 ± 2.69a | 103.06 ± 3.19a | 8.39 ± 0.19a |
| *O. glaberrima* (IR 102925) | Cultivated | 30.41 ± 0.14c | 70.04 ± 0.79cd | 149.21 ± 1.80c | 10.28 ± 0.10c |
| *O. latifolia* (IRGC 99596) | Wild | 34.32 ± 0.39d | 76.75 ± 1.24de | 166.17 ± 2.86d | 11.72 ± 0.27d |
| *O. australiensis* (IRGC 105272) | Wild | 36.81 ± 0.37e | 81.25 ± 1.08e | 181.93 ± 0.77e | 12.27 ± 0.12d |

Next, we calculated the photosynthetic limitations due to *V_cmax_* (*A_c_*), *J_max_* (*A_j_*), and *TPU* (*A_t_*). The transition from Rubisco-limited (*A_c_*) to RuBP-regeneration limited (*A_j_*) photosynthesis in Asian cultivated rice, IR 64 and Nipponbare, was observed at *C_i_* < 250 µmol mol^-1^ (Supplementary Table S5). In contrast, wild rice species *O. australiensis* and *O. latifolia* along with African cultivated rice *O. glaberrima* showed *A_c_* to *A_j_* transition at *C_i_* > 300 µmol mol^-1^. Similarly, the transition from RuBP-regeneration limited (*A_j_*) to TPU-limited (*A_t_*) photosynthesis in Nipponbare and IR 64 was observed at *C_i_* 524 and 608 µmol mol^-1^, respectively, which was lesser compared to *O. glaberrima* (657 µmol mol^-1^), *O. australiensis* (700 µmol mol^-1^), and *O. latifolia* (713 µmol mol^-1^) (Supplementary Table S5). CO_2_-saturated *A_max_* that ranged from 21.37 µmol CO_2_ m^-2^ s^-1^ in IR 64 to 36.81 µmol CO_2_ m^-2^ s^-1^ in *O. australiensis*, further, indicated TPU-limitation (*A_t_*) in IR64 and Nipponbare compared with the wild rice accessions. However, the TPU-limitation is observed at *C_i_* much higher than the physiological conditions as reported earlier (Kumarathunge *et al*., 2019). These results suggested that leaf photosynthesis per unit area in Asian cultivated varieties, IR 64 and Nipponbare, was limited due to Rubisco activity (*V_cmax_*) and RuBP-regeneration/Electron Transport Rate (*J_max_*) compared to the selected wild rice genotypes under ambient CO_2_ concentrations.

### Photochemical differences among the selected rice accessions

Since the estimation of biochemical limitations to photosynthetic differences across the selected accessions suggested *J_max_* as one of the limiting factors, we used leaf chlorophyll fluorescence parameters to quantify the photochemical features of the selected rice accessions. We observed strong differences in electron transport rate (*J_PSII_*), the efficiency of PSII (□*PSII*), and photochemical quenching (*qP*) among the selected cultivated and wild rice accessions across different CO_2_ concentrations, with distinct differences at CO_2_ concentration higher than 200 µmol mol^-1^. *O. latifolia* and *O. australiensis* showed higher values of *J_PSII_* than the cultivated Asian rice varieties IR 64 and Nipponbare at different CO_2_ concentrations, with 19% to 53% higher *J_max_* in the two wild rice accession compared with the two cultivated varieties (Fig. 2A). Similar to *J_PSII_*, values of □*PSII* and *qP* were also significantly higher for *O. latifolia* and *O. australiensis* compared with the cultivated varieties (Fig. 2B, C). *O. glaberrima*, the African cultivated rice with reasonably higher *A*, also showed relatively higher *J_PSII_*, □*PSII*, and *qP* than *O. sativa* cultivated varieties at different CO_2_ concentrations. Similar response patterns of both *A* and *J_PSII_* were observed at different CO_2_ concentrations, and were found to be strongly correlated with each other for all the selected genotypes (Supplementary Fig. S8). Taken together, these results showed that photochemical features of *O. australiensis* and *O. latifolia* contributed to the higher leaf photosynthesis rate, and lower *J_PSII_*, □*PSII*, and *qP* of IR 64 and Nipponbare accounted for RuBP-regeneration-limited photosynthesis in those cultivated varieties.

**Fig. 2.**
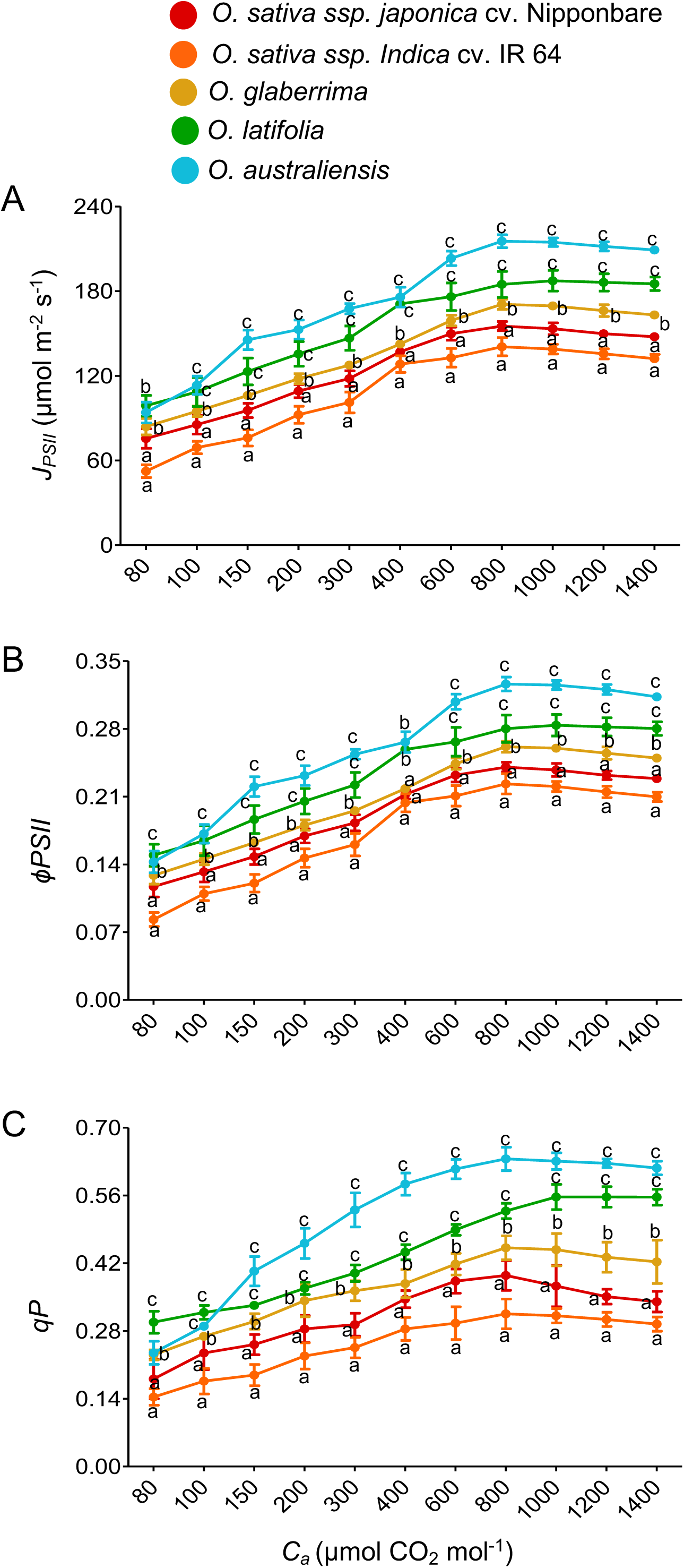
Photochemical features of the selected cultivated and wild rice accessions. Quantification of electron transport rate, *J_PSII_* (A), the efficiency of PSII, □*PSII* (B), and photochemical quenching, *qP* (C) of the selected accessions at varying CO_2_ concentrations. Each value represents mean ± SE, where n = five data points from different plants. Different letters at each CO_2_ concentration indicate statistical significance according to one-way ANOVA followed by post-hoc Tukey HSD calculation at *P ≤ 0.05*.

### Rubisco activity differences across the selected rice genotypes

Our analyses on biochemical limitations to photosynthesis suggested *V_cmax_* as a major limitation to leaf photosynthesis per unit area in the cultivated varieties IR64 and Nipponbare. Therefore, we quantified the *RuBP-use* and *in vitro* Rubisco activity (*V_cmax_*) over 30 minutes across selected cultivated and wild rice species (Fig. 3A, B). The CO_2_-saturated *RuBP-use* gradually increased immediately from 10 sec to 10 min, followed by saturation for all the accessions. The *RuBP-use* was significantly higher in photosynthetically efficient wild rice accessions (*O. latifolia* and *O. australiensis*) than that of Asian cultivated varieties (*O. sativa* cv. IR 64, and *O. sativa* cv. Nipponbare) (Fig. 3A). Consistent with the *RuBP-use*, V*_cmax_* was maximum for the wild rice *O. australiensis* and minimum for the cultivated variety IR64 at saturation (Fig. 3B). Table 4 shows significantly higher values of *RuBP-use* per sec and *V_cmax_* for *O. latifolia* and *O. australiensis* compared to IR 64 and Nipponbare at different time points during the assay. The results not only supported the estimated photosynthetic limitations due to *V_cmax_* in the selected cultivated varieties but also suggested the existence of variation in Rubisco carboxylation activity per unit leaf area within the rice system.

**Fig. 3.**
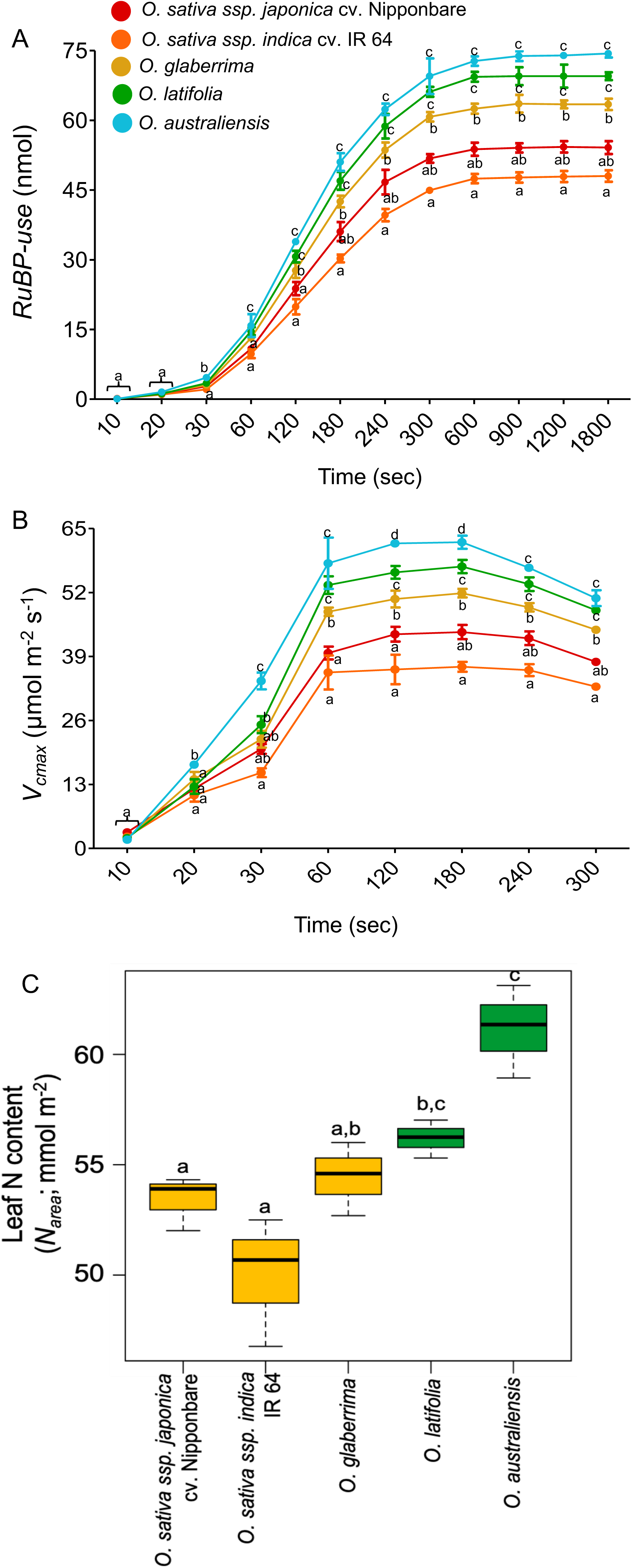
RuBP-use, Rubisco activity, and leaf nitrogen content across the selected accessions. (A) NADH-linked spectrophotometric measurement of RuBP-use (nmol) at different time points from 10 to 1800 sec. (B) Quantification of *V_cmax_* for the selected accessions at different time points up to saturation. (C) Leaf nitrogen content/unit leaf area (N*_area_*) in the flag leaves of the selected accessions. Values represent mean ± SE (three data points from different plants). Different letters indicate statistical significance according to one-way ANOVA followed by post-hoc Tukey HSD calculation at *P* ≤ *0.05*.

**Fig. 4.**
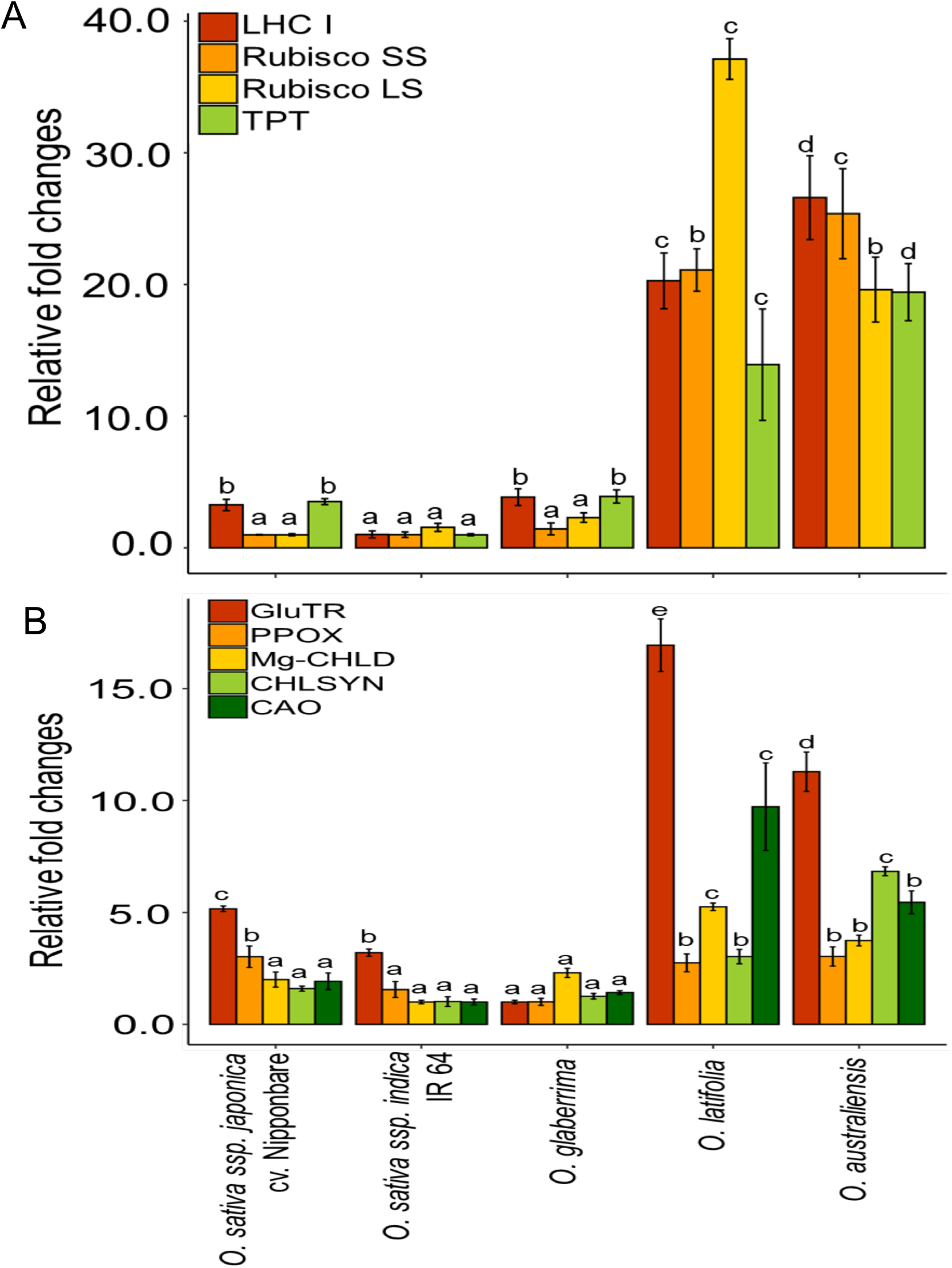
Expression pattern of photosynthetic and chlorophyll biosynthetic genes across the selected accessions. (A) Expression levels of genes encoding a key component of the light-harvesting complex, Rubisco, and Triose phosphate/phosphate translocator. (B) Expression levels of key chlorophyll biosynthetic genes. Relative expressions of the target genes are normalized using rice actin (LOC_Os03g50885.1) as the internal control. Values represent mean ± SE (three data points from different plants). Different letters indicate statistical significance according to one-way ANOVA followed by post-hoc Tukey HSD calculation at *P* ≤ *0.05*. Gene abbreviations: LHCI, Light-harvesting chlorophyll protein complex I; Rubisco SS, Ribulose bisphosphate carboxylase small subunit; Rubisco LS, Ribulose bisphosphate carboxylase large subunit; TPT, Triose phosphate/phosphate translocator; GluTR, Glutamyl-tRNA reductase; PPOX, Protoporphyrinogen oxidase; Mg-CHLD subunit ChlD, Magnesium chelatase; CHLSYN, Chlorophyll synthase; CAO, Chlorophyllide a oxygenase.

**Table 4.** *in vitro* RuBP-use and Rubisco carboxylation rate (*V_cmax_*) for selected rice genotypes. Each value represents mean ± SE, where n = three data points from different plants. The significant difference among the accessions is indicated by different letters (one-way ANOVA followed by Tukey’s HSD, *P* < 0.05).

| Genotypes | Cultivated/<br>Wild | *RuBP-use* (nmol s<sup>-1</sup>) | | | $V_{cmax}$ (μmol m<sup>-2</sup> s<sup>-1</sup>) | | |
| --- | --- | --- | --- | --- | --- | --- | --- |
|  |  | 120 s | 180 s | 240 s | 120 s | 180 s | 240 s |
| *O. sativa* cv. Nipponbare | Cultivated | 0.198 ± 0.01b | 0.200 ± 0.01b | 0.195 ± 0.01b | 43.52 ± 1.50b | 43.98 ± 1.43b | 42.71 ± 1.40b |
| *O. sativa* cv. IR 64 | Cultivated | 0.166 ± 0.01a | 0.168 ± 0.01a | 0.165 ± 0.01a | 36.37 ± 3.00a | 36.91 ± 1.00a | 36.22 ± 1.22a |
| *O. glaberrima* (IR 102925) | Cultivated | 0.231 ± 0.01cd | 0.236 ± 0.01c | 0.223 ± 0.01c | 50.67 ± 1.71c | 51.87 ± 0.87c | 49.01 ± 0.85c |
| *O. latifolia* (IRGC 99596) | Wild | 0.256 ± 0.01d | 0.261 ± 0.01d | 0.245 ± 0.01cd | 56.10 ± 1.30d | 57.29 ± 1.35d | 53.69 ± 1.39d |
| *O. australiensis* (IRGC 105272) | Wild | 0.282 ± 0.01d | 0.284 ± 0.01d | 0.260 ± 0.01d | 61.99 ± 0.48d | 62.22 ± 1.35d | 57.02 ± 0.65d |

The differences in Rubisco activity per unit leaf area across the selected accessions may be attributed either to Rubisco amount or to enzyme kinetics differences. Since the leaf nitrogen content would inform about Rubisco amount per unit leaf area, we quantified the leaf nitrogen content of the selected accessions. Photosynthetically efficient *O. australiensis* and *O. latifolia* accessions showed significantly higher nitrogen content per unit leaf area (*N_area_*) compared with IR64 and Nipponbare (Fig. 3C). The selected *O. australiensis* accession had 22% and 14% higher *N_area_*, whereas *O. latifolia* accession had 12% and 5% higher *N_area_* than IR64 and Nipponbare, respectively, suggesting an important contribution of leaf nitrogen content to higher Rubisco activity per unit area in the selected wild accessions.

### Expression pattern of genes encoding core photosynthetic and chlorophyll biosynthesis enzymes

Since the differences in the Rubisco amount per unit leaf area may also contribute to the Rubisco-limitation of the photosynthesis in IR 64 and Nipponbare, we checked the expression pattern genes encoding Rubisco small and large subunits. In addition, we checked the expression of LHC1 (Light-harvesting chlorophyll protein complex) and TPT (Triose phosphate translocator) that may contribute to RuBP-regeneration- and TPU-limitations of the photosynthesis, respectively (Supplementary Table S3). The expression of the selected photosynthetic genes was significantly higher in *O. latifolia* and *O. australiensis* compared to the selected cultivated varieties (Fig. 4A, Supplementary Fig. S9A), suggesting higher photosynthesis in those species could, at least in part, be due to the higher abundance of these gene products. These genes could increase net photosynthetic efficiency by increasing the efficiency of light capture, CO_2_ assimilation, and transport of triose phosphate from chloroplast to cytoplasm. Since wild rice species with higher photosynthesis also showed higher chlorophyll content, we checked the expression level of chlorophyll biosynthesis genes. The expression of chlorophyll biosynthesis genes, such as *GluTR*, *PPOX*, *Mg-CHLD*, *CHLSYN*, and *CAO* were found to be significantly higher in *O. latifolia* and *O. australiensis* compared to the selected cultivated varieties (Fig. 4B, Supplementary Fig. S9B).

### Leaf developmental traits underpinning photosynthetic variations

Leaves are the prime photosynthetic organs, and expectedly the selected cultivated and wild rice accessions exhibited considerable variation for flag leaf width (FLW) and thickness (FLT) (Supplementary Fig. S10). Wild rice species had thicker and wider leaves as compared to cultivated rice. Strong significant positive correlations of leaf photosynthesis were found with FLW (*r = 0.850*) and FLT (*r = 0.938)* (Fig. 5A). At the anatomical level, remarkable differences in the vascular architecture and mesophyll cell features were observed for the selected rice accessions that differ in leaf photosynthesis (Supplementary Table S6). Wild rice species with higher *A* exhibited a larger vein height and width. *O. sativa* cv. Nipponbare that showed relatively higher *A* compared to other cultivated varieties also showed larger vein height and width (Supplementary Table S6). Significant positive correlations were observed for *A* with vein features: minor vein width (MiV-W, *r = 0.752*), minor vein height (MiV-H, *r = 0.899*), and total number of veins (TV, *r = 0.848*) (Fig. 5A). Similar strong correlations of *A* were observed for mesophyll cell features: mesophyll cell lobbing (M-Lo, *r = 0.760*), mesophyll cell length (M-L, *r = 0.863*), mesophyll cell width (M-W, *r =0.928*), and mesophyll cell area (M-A, *r = 0.897*) at *P* ≤ 0.05. However, no significant correlations of *A* and related physiological traits were observed with bundle sheath features: bundle-sheath number (BS-No), and bundle-sheath area (BS-A) (Fig. 5A).

**Fig. 5.**
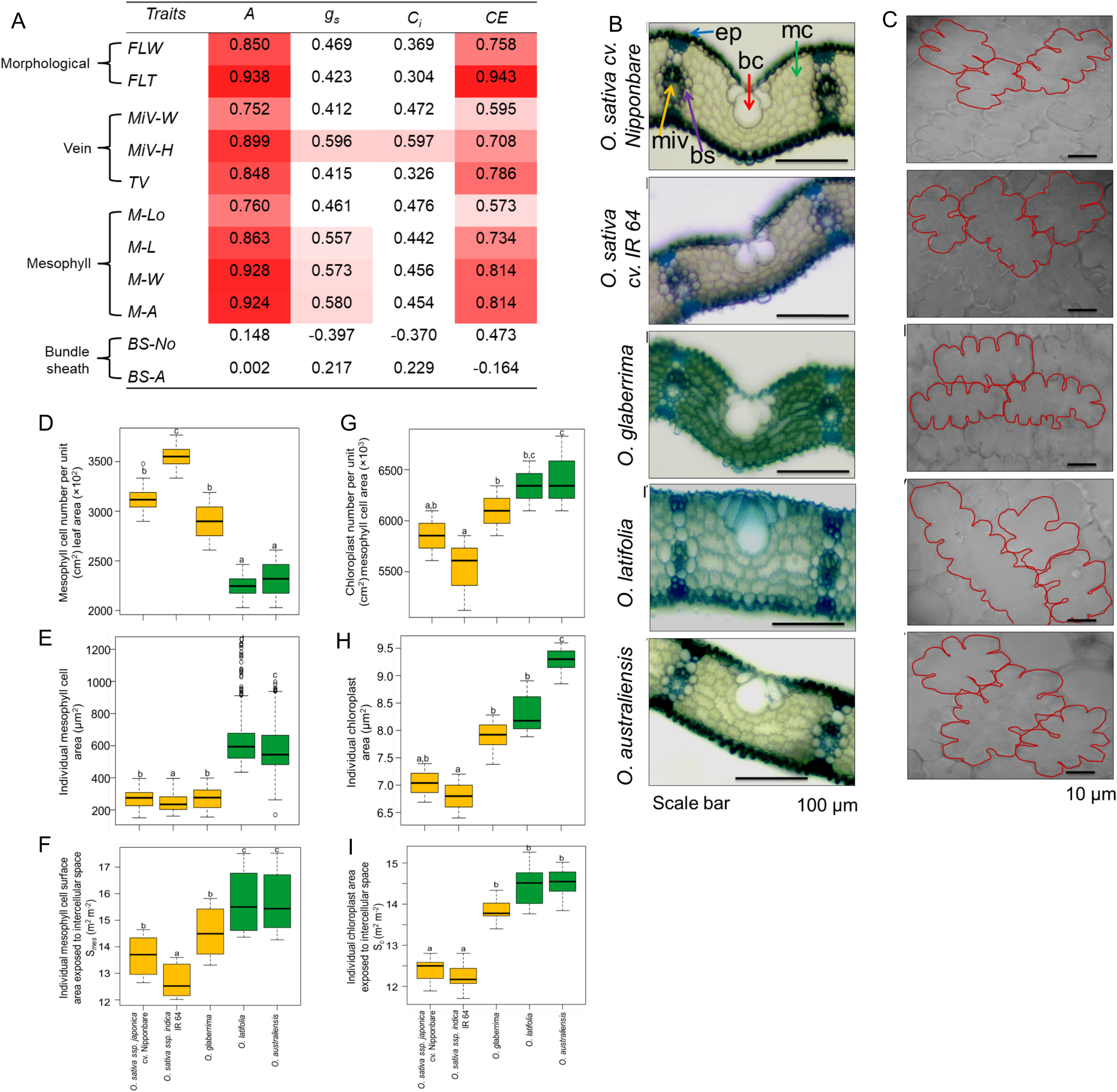
Leaf developmental traits affecting photosynthesis rate and related physiological traits of rice. (A) Correlation of leaf photosynthesis rate and related physiological traits with leaf developmental features. The red shade represents a significant positive correlation at *P* ≤ *0.05*. (B) Bright-field images of a cross-section of a flag leaf of the selected genotypes showing variations in mesophyll cells and vasculature. Arrows indicate bulliform cells (bc), bundle sheath (bs), epidermis (ep), mesophyll cells (mc), and minor veins (miv). Scale bars represent 100 µm. (C) Bright-field microscopy images showing variations in mesophyll cell features, including lobbing. Scale bar represents 10 µm. (D-F) Quantification of mesophyll cell number per unit (cm^2^) leaf area (D), individual mesophyll cell area (E), and individual mesophyll cell surface area exposed to intercellular space (*S_mes_*, F) of the selected cultivated and wild rice accessions. (G-I) Quantification of chloroplast number per unit (cm^2^) mesophyll cell area (G), individual chloroplast area (H), and individual chloroplast area exposed to intercellular space (*S_c_*, I) of the selected cultivated and wild rice accessions. Cultivated rice accessions and wild species are represented with yellow- and green-shaded boxes, respectively. Each box plot shows the median and interquartile range with minimum and maximum values. Different letters indicate statistical significance according to one-way ANOVA followed by post-hoc Tukey HSD calculation at *P* ≤ *0.05*.

Photosynthetically efficient wild rice species, *O. latifolia,* and *O. australiensis*, had larger and closely spaced veins with a fewer number of mesophyll cells between two consecutive veins (Fig. 5B). Those wild species also had larger mesophyll cell size, and thus a lesser number of mesophyll cells per unit leaf area (Fig. 5C, D). We, then, quantified attributes related to the mesophyll cell surface area of the selected accessions. Wild rice accessions of *O. australiensis* and *O. latifolia*, showed remarkably higher average mesophyll cell area (2.2 to 2.6-fold larger mesophyll cells) as well as significantly higher mesophyll cell surface area exposed to intercellular space (*S_mes_*) compared with IR64 and Nipponbare (Fig. 5E, F). We also investigated the organization of chloroplasts in mesophyll cells of the representative wild and cultivated accessions. Photosynthetically efficient wild rice exhibited larger and numerous chloroplasts, which were systematically distributed along the mesophyll cell walls (Fig. 5G, H; Supplementary Fig. S11). The surface area of chloroplasts exposed to intercellular space (*S_c_*) was also found to be significantly higher in the photosynthetically efficient wild rice accessions compared with the cultivated varieties IR64 and Nipponbare (Fig. 5I). These results underline the strong influence of mesophyll and chloroplast features in driving high leaf photosynthesis per unit area in the selected wild rice accessions.

## Discussion

The importance of investigating variations in leaf photosynthesis within and across species to be exploited for crop improvement has been emphasized ( Ort *et al*., 2015; Adachi *et al*. 2019; Simkin *et al*., 2019; van Bezouw *et al*., 2019). Exploiting the photosynthetic variations within the rice system itself, including cultivated and wild rice, could be one of the potential strategies to increase genetic yield potential. In this study, we not only quantified leaf photosynthetic differences across the selected cultivated and wild rice accessions but also identified the biochemical limitations to leaf photosynthesis per unit area in the selected cultivated varieties, IR64 and Nipponbare. In addition, we also identified photochemical and leaf developmental traits associated with higher leaf photosynthesis rates in the selected wild rice accessions of *O. australiensis* and *O. latifolia*.

### High leaf photosynthesis rate in the selected wild rice accessions, and related physiological traits

Wild relatives of rice are shown to have more genetic variation compared with cultivated varieties as domestication was primarily targeted for optimal plant architectural and grain features (Huang *et al*., 2012; Chen *et al*., 2021). Therefore, the wild relatives are suggested to have useful variants for various desirable agronomic traits. Indeed, wild relatives of rice have been used for breeding stress tolerance in cultivated rice varieties (Sanchez *et al.,* 2013; Atwell *et al*., 2014; Wing *et al*., 2018). We found higher leaf photosynthesis per unit area (*A*) in the selected accessions of wild rice *O. australiensis*, *O. latifolia*, and *O. rufipogon*, as well as African domesticated rice *O. glaberrima* compared with the cultivated varieties of *Oryza sativa* (Fig. 1A). A few previous studies have reported high leaf photosynthesis rate in some of the wild relatives of rice, including *O. australiensis* and *O. rufipogon*, than cultivated species *O. sativa* (Zhao *et al*., 2010; Kiran *et al*., 2013; Kondamudi *et al.,* 2016; Haritha *et al.,* 2017). However, a few other studies found no obvious difference or lower *A* in wild relatives of rice compared with *O. sativa*, suggesting that high *A* is not a general feature of wild rice relatives, rather an accession-specific feature (Yeo *et al.,* 1994; Scafaro *et al*., 2011; Giuliani *et al*., 2013). Higher leaf photosynthesis rate could be attributed to differences in the life strategies of the selected wild rice accessions compared with the cultivated varieties. The perennial nature of the wild accessions with remarkably higher biomass accumulation than cultivated varieties might require more photosynthates, and thus increasing leaf photosynthesis rate. We have recently shown preferential utilization of a higher amount of photosynthates in the vegetative tissues of the specific *O. australiensis* accession for the synthesis of structural carbohydrates (Mathan *et al*., 2021). Furthermore, a higher amount of photosynthates would also support more organ initiation and larger organ size, as well as the general maintenance of the higher biomass in the selected wild rice accessions. Thus, the requirement of photosynthates for general growth and biomass as well as specific biochemical, photochemical, and developmental features would determine the accession-specific value of *A*. Therefore, understanding the underlying traits that drive high *A* in specific accessions is crucial.

Consistent with leaf photosynthesis rate, photosynthetically efficient wild rice accessions of *O. australiensis* and *O. latifolia* showed higher values of stomatal conductance (*g_s_*) and mesophyll conductance (*g_m_*) that would facilitate higher CO_2_ concentration in intercellular spaces (*C_i_*) as well as inside mesophyll cells of those species (Fig. 1B, C; Table 2). Further, the wild rice accession also had reasonably higher *CE* than cultivated varieties. Higher leaf nitrogen content, indicating more Rubisco per unit leaf area, supports higher *CE* and *A* in the selected wild rice accessions than the cultivated varieties (Fig. 3C). The *A* has been shown to be positively associated with *CE* in rice (Yeo *et al*., 1994). Indeed, quantification of photosynthesis and related physiological parameters for introgressions from *O. rufipogon* in a cultivated background showed the influence of both *CE* as well as *g_s_* on higher leaf photosynthesis (Haritha *et al*., 2017). Comparison of several cultivated and wild rice genotypes had shown a close positive relation between *A* and *g_m_*, and *g_m_* could increase leaf photosynthesis per unit area without affecting transpiration (Flexas *et al*., 2008; Zhu *et al*., 2010; Giuliani *et al*., 2013). Therefore, efforts to increase *g_s_*, *g_m_*, and *CE* in the cultivated rice varieties comparable to the high photosynthetic wild accessions will help in increasing photosynthetic efficiency.

### Rubisco carboxylation rate (*V_cmax_*) and electron transport rate (*J_max_*) promote leaf photosynthesis rate in the selected wild rice accessions

Rubisco is the rate-limiting enzyme for photoassimilation of CO_2_, and Rubisco abundance and activity strongly determine the photosynthetic performance of a plant (Parry *et al*., 2013; Sage, 2002). Higher Rubisco activity per unit leaf area in the selected wild accessions compared with the cultivated varieties could be due to either more Rubisco per unit area or the presence of an efficient Rubisco in the wild species. High photosynthetic wild accessions had higher expression of genes encoding Rubisco large subunit and small subunit, indicating the more abundance of Rubisco in those species (Fig. 4A). Thick leaves of the photosynthetically efficient wild species would support the first alternative where increasing leaf thickness would lead to more Rubisco per unit area. More Rubisco per unit area, in addition, could also be achieved by more nitrogen allocation to Rubisco. Higher nitrogen content per unit area in the leaves of the photosynthetically efficient *O. australiensis* and *O. latifolia* accessions than the cultivated varieties would also support higher Rubisco content per unit area in the wild accession (Fig. 3C). Therefore, possible ways to increase Rubisco content per unit leaf area may overcome the Rubisco limitation of leaf photosynthesis in the cultivated varieties. It would, further, be interesting to investigate if there are differences in Rubisco kinetic properties across photosynthetically contrasting cultivated and wild rice that might contribute to differences in Rubisco carboxylation activity.

The comparative higher values for the quantum efficiency of PSII ( □*PSII*) and photochemical quenching (*qP*) along with *J_PSII_* in *O. australiensis* and *O. latifolia* accessions indicate more receptive PSII reaction centres for electron transport from PSII to PSI in those species, making an efficient link between light and light-independent reactions to increase *A* (Fig. 2A-C) (Kramer and Evans, 2011). Gene expression studies also showed higher expression of the *LHC1* gene, which is involved in the organization of reaction centre, in photosynthetically efficient wild species. Results, further, showed higher expression of chlorophyll biosynthetic genes and higher chlorophyll content in those wild species, aiding the efficient light harvesting (Fig. 1E, 4B). Understanding the differential regulation of genes encoding components of photosystem as well as chlorophyll biosynthesis in the wild species could help in improving photochemical features in the cultivated varieties.

### Targeting leaf thickness and width to overcome the biochemical limitations to leaf photosynthesis in the cultivated varieties

We observed a strong positive association of *A* with leaf width and thickness (Fig. 5A). Leaf thickness has been suggested to influence photosynthesis through efficient absorption and utilization of the light by homogenizing the leaf internal light distribution as well as maintaining leaf temperature (Tholen *et al*., 2012). Increasing leaf thickness, and thus more tissue per unit area would be instrumental for more nitrogen and Rubisco content, chlorophyll content, and electron transport machinery per unit area. Higher chlorophyll and electron transport machinery per unit area would facilitate efficient light harvesting for RuBP regeneration, whereas more Rubisco content per unit leaf area would increase carboxylation efficiency. Thus, targeting leaf thickness, similar to the photosynthetically efficient wild rice, may result in high net photosynthesis per unit leaf area in the cultivated varieties by overcoming Rubisco and electron transport rate limitations. Such leaves would indeed be beneficial under high light intensity, but may not be optimal for light harvesting and photosynthesis under low light conditions. Nonetheless, increasing leaf thickness, resulting in higher nitrogen and Rubisco content per unit area, could be a potential way to overcome the biochemical limitation of photosynthesis in the cultivated rice varieties under ambient light conditions. The association of leaf width with photosynthetic performance and yield in rice has been observed earlier. *WUSCHEL-RELATED HOMEOBOX* (*WOX*) genes *NAL1*, *NAL2*, and *NAL3* are associated with increased leaf width, which in turn results in increased photosynthetic efficiency and yield (Fujita *et al*., 2013; Ishiwata *et al*., 2013). Introgression of a natural variant of *NAL1* from *japonica* to *indica* cultivar led to thicker and wider leaves with higher leaf photosynthesis rate and yield (Takai *et al*., 2013).

### Distinct leaf anatomical features of the photosynthetically efficient wild rice accessions

Photosynthetically efficient wild rice accessions have mesophyll cell features that promote both higher accesses to available CO_2_ as well as efficient light harvesting. Larger sized and lobed mesophyll cells along with more surface area exposed to intercellular spaces in wild rice accessions likely provide increased surface area to access more internal CO_2_ (Fig. 5C-F), whereas the higher number of larger and systematically distributed chloroplasts in the mesophyll cells of those species may promote efficient light capture to enhance photosynthesis (Fig. 5G, H; S11). In addition, higher values of *S_mes_* and *S_c_* would support the higher *g_m_* of the photosynthetically efficient wild rice accessions, leading to efficient CO_2_ diffusion inside the mesophyll cells (Adachi *et al*. 2013; Pathare *et al*., 2020). These, together with thicker leaves having more nitrogen and Rubisco content per unit area, would provide a higher amount of photosynthetic inputs as well as machinery in the photosynthetically efficient wild rice accessions. Lesser number of mesophyll cells between two consecutive veins in photosynthetically efficient wild species, a feature similar to C_4_ plants, would be promoting efficient transport of photoassimilate (Amiard *et al*., 2005; Feldman *et al*., 2017). Larger vein size in the wild rice species would, further, be adding to the efficient transport of photoassimilates, thus increasing the leaf photosynthesis (Sack and Scoffoni, 2013). Thus, photosynthetically efficient wild rice accessions with desirable anatomical features provide a potential genetic resource towards the identification of genes for desirable manipulation in mesophyll, vein, and chloroplast features.

In summary, we identified biochemical limitations to leaf photosynthesis per unit area in the selected cultivated rice varieties, IR64 and Nipponbare, compared to photosynthetically efficient wild rice accessions of *O. australiensis* and *O. latifolia*. Rubisco carboxylation activity and electron transport rate along with photochemical features (quantum efficiency of PSII and Photochemical quenching) and leaf developmental features (thicker and wider leaves, larger and lobbed mesophyll cells, larger veins, and lesser number of mesophyll cells between two consecutive veins) contribute to higher leaf photosynthesis in the selected *O. australiensis* and *O. latifolia* accessions (Fig. 6). The distinct leaf anatomy in the selected wild accessions would provide structure to accommodate more nitrogen and photosynthetic machinery per unit leaf area than the cultivated varieties. Similarly, higher *S_mes_* and *S_c_* would lead to more CO_2_ diffusion inside the mesophyll cells and chloroplasts of those wild accessions. Together, these anatomical features would mediate higher Rubisco activity and electron transport rate driving higher leaf photosynthesis rate in the selected wild accessions. Future efforts to dissect the genetic and molecular basis of the underlying biochemical, photochemical, and developmental traits shall be instrumental in tailoring photosynthetically efficient rice towards the possible increases in genetic yield potential.

**Fig. 6.**
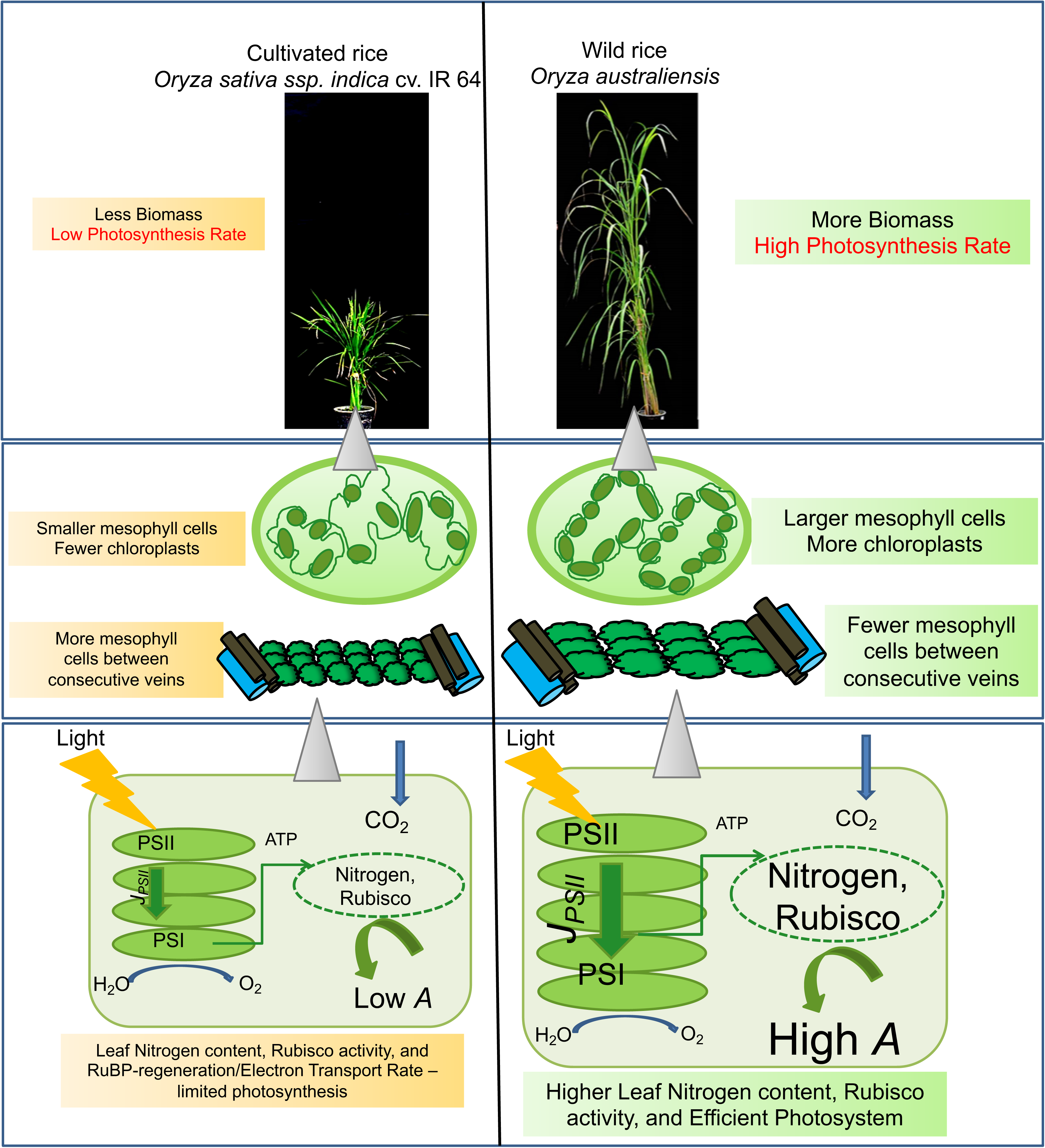
Leaf developmental, photochemical, and biochemical traits contributing towards photosynthetic differences between two extreme genotypes: *Oryza sativa* ssp. *indica* cv. IR 64 and *Oryza australiensis*. Wider leaf, larger mesophyll cells, larger veins, fewer number of mesophyll cells between two consecutive veins, more chloroplasts, efficient Electron Transport Rate along with higher leaf nitrogen content and Rubisco activity facilitate higher leaf photosynthesis per unit area in the wild rice species *Oryza australiensis* compared to the cultivated variety *Oryza sativa* ssp. *indica* cv. IR 64.

## Supplementary data

**Fig. S1.** Pair-wise Pearson correlation coefficient (*r*) analysis among the leaf photosynthesis and physiological traits.

**Fig. S2.** Representative images of growth-chambers-grown plants of the selected cultivated and wild rice accessions.

**Fig. S3.** Effect of varying light intensity and CO_2_ concentration on leaf photosynthesis and stomatal conductance.

**Fig. S4.** *C_i_* values for light-response curves and CO_2_-response curves for each genotype.

**Fig. S5.** Representative graphs for the quantification of mitochondrial respiration rate in the light (*R*_d_) and the CO_2_ compensation point related to *C_i_* (Γ*\**) for each genotype.

**Fig. S6.** Relationship between *A* and *C_i_* with *A*/*C_i_* curve fitting for the selected cultivated and wild rice accessions.

**Fig. S7.** Linear regression analysis between *V_cmax_* and *J_max_* (A), *TPU* and *J_max_* (B), and *V_cmax_* and *TPU* (C).

**Fig. S8.** Response and correlation of leaf photosynthesis (*A*) and electron transport rate (*J_PSII_*) at varying CO_2_ concentrations for the selected rice accessions.

**Fig. S9.** Expression pattern of photosynthetic and chlorophyll biosynthetic genes across the selected accessions normalized to rice ubiquitin internal control.

**Fig. S10.** Flag leaf width and thickness of the selected cultivated and wild rice accessions.

**Fig. S11.** TEM images for the organization of chloroplasts in mesophyll cells.

**Table S1.** Details of rice accessions used in the present study.

**Table S2.** Total chlorophyll content (*TCC*, µmol m^-2^) and leaf absorbance (α) of fully-grown eighth leaves of plants grown under controlled conditions.

**Table S3.** Details of genes used for expression analysis.

**Table S4.** List of primers for qRT-PCR analysis.

**Table S5.** *A_c_*-*A_j_* and *A_j_*-*A_t_* transitions for the selected genotypes.

**Table S6.** Quantification of leaf developmental traits of the fully expanded flag leaves of the selected cultivated rice varieties and wild rice species.

## Author contributions

JM and AR conceptualized the study. JM, AS, VJ, and AR designed experiments. JM, AS, and VJ performed experiments, analysed data, and compiled figures. AS and AR wrote the manuscript. All authors contributed to and edited the final manuscript.

## Data availability statement

Data sharing is not applicable to this article as all created data is already contained within this article or in the supplementary material.

## Supporting information

Supplemental Fgure and Table

## Acknowledgments

This work was supported by the core funding from the National Institute of Plant Genome Research as well as Ramalingaswamy Re-entry Fellowship (BT/RLF/Re-entry/05/2013) and Innovative Young Biotechnologist Award (BT/09/IYBA/2015/01) from the Department of Biotechnology, Ministry of Science and Technology, India. JM, AS, and VJ acknowledge their CSIR-JRF, SERB-NPDF, and UGC-JRF, fellowships, respectively. Seeds of wild rice species and *Oryza glaberrima* were kindly provided by Dr. Kuldeep Singh and Dr. Kumari Neelam, Punjab Agricultural University, Ludhiana, India. We thank Dr. Subodh K Sinha for his help in leaf nitrogen content estimation.

## Notes

### Competing Interest Statement

The authors have declared no competing interest.

