## Supplemental Fgure and Table for "High photosynthesis rate in the selected wild rice is driven by leaf anatomy mediating high Rubisco activity and electron transport rate"

| *Traits* | *A* | $g_s$ | $C_i$ | *CE* |
|---|---|---|---|---|
| $g_s$ | 0.629 | | | |
| $C_i$ | 0.570 | 0.965 | | |
| *CE* | 0.819 | 0.102 | 0.002 | |
| *TCC* | 0.912 | 0.360 | 0.310 | 0.906 |

**Fig. S1.** Pair-wise Pearson correlation coefficient (*r*) analysis among the leaf photosynthesis and physiological traits. The red shade represents the significant positive correlation coefficient at $P \leq 0.05$.

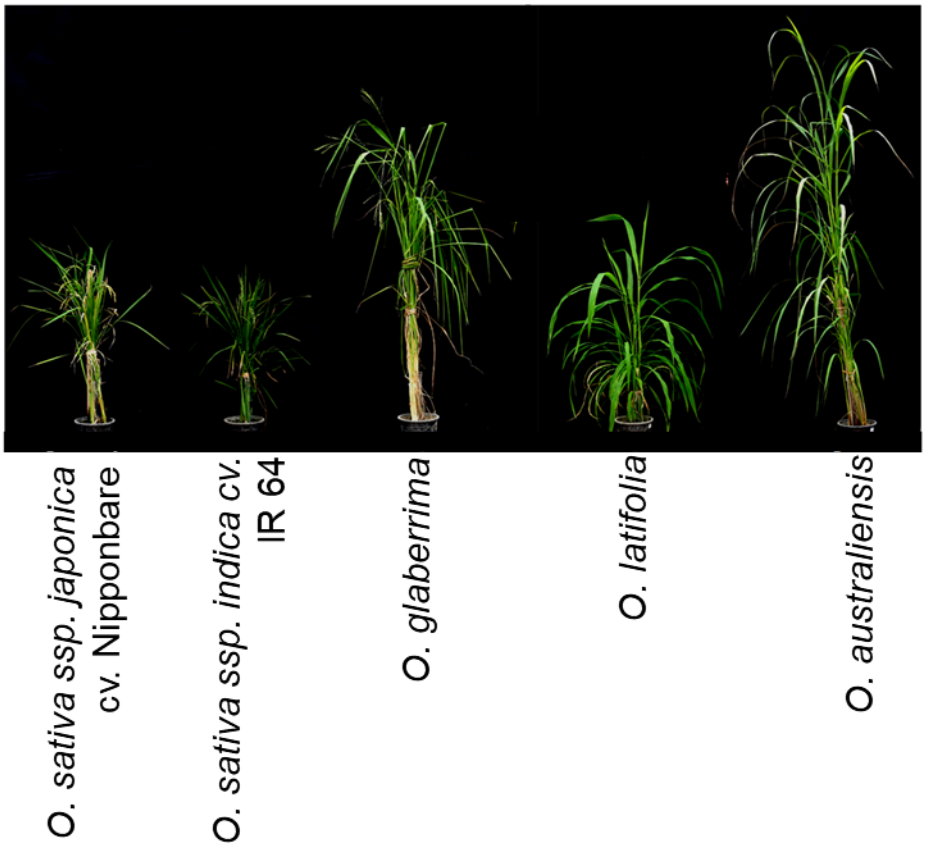


**Fig. S2.** Representative images of controlled growth-chambers-grown plants of the selected cultivated and wild rice accessions.

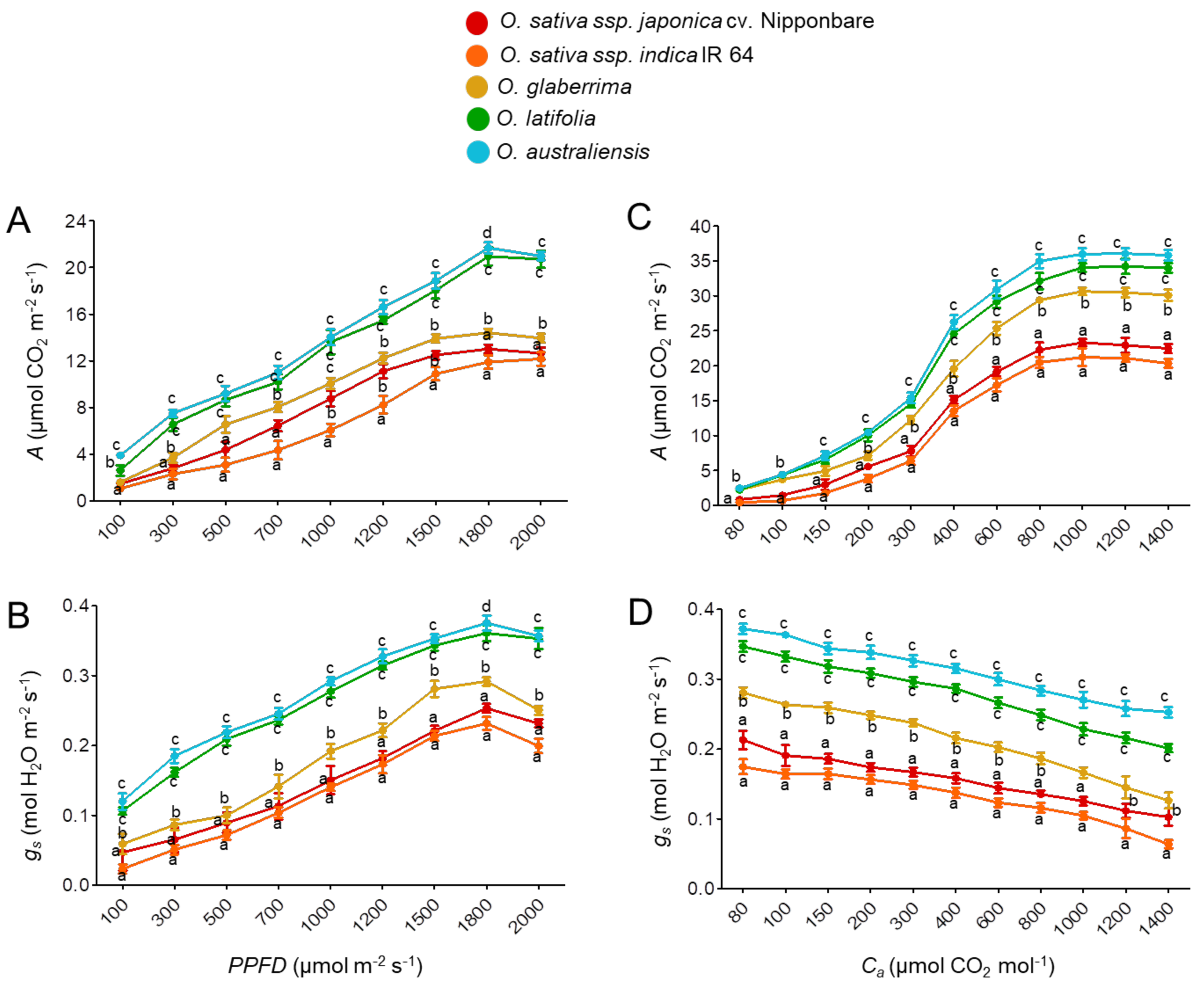


**Fig. S3.** Effect of varying light intensity and $CO_2$ concentration on leaf photosynthesis and stomatal conductance.

(A-B) Quantification of net photosynthesis per unit leaf area, *A* (A) and stomatal conductance to water, $g_s$ (B) in response to increasing PPFD from 100 to 2000 µmol m$^2$ s$^{-1}$ at 400 µmol mol$^{-1}$ $CO_2$ concentration.

(C-D) Quantification of net photosynthesis per unit leaf area, *A* (C) and stomatal conductance to water, $g_s$ (D) in response to varying external $CO_2$ concentration ($C_a$) at 1500 µmol m$^{-2}$ s$^{-1}$ light intensity.

Each value represents mean ± SE, where n = five data points from different plants. Different letters at each light intensity and $CO_2$ concentration indicate statistical significance according to one-way ANOVA followed by post-hoc Tukey HSD calculation at $P \leq 0.05$.

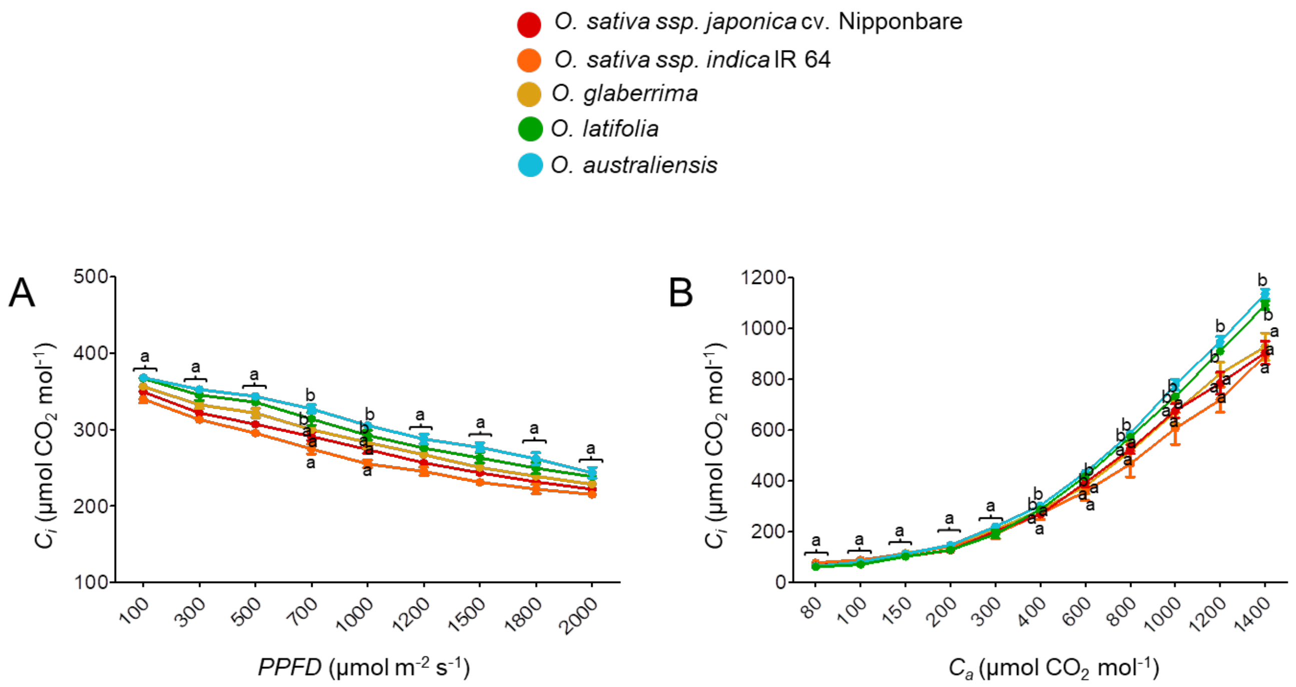


**Fig. S4.** $C_i$ values for light-response curve (A) and $CO_2$-response curve (B) for each genotype. Each value represents mean ± SE, where n = five data points from different plants. Different letters at each light intensity and $CO_2$ concentration indicate statistical significance according to one-way ANOVA followed by post-hoc Tukey HSD calculation at $P \leq 0.05$.

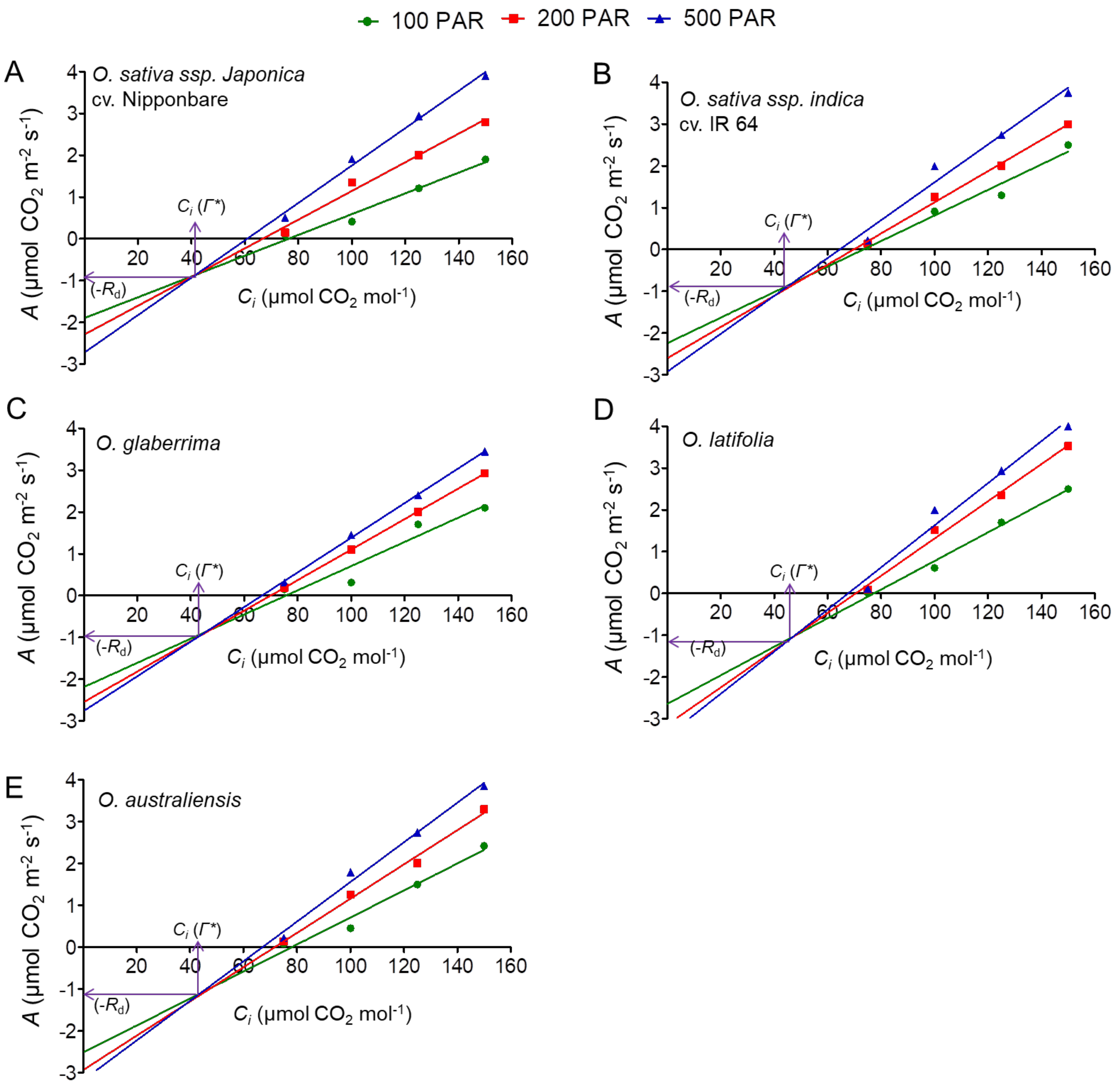


**Fig. S5.** Representative graphs for the quantification of mitochondrial respiration rate in the light ($R_d$) and the $CO_2$ compensation point related to $C_i$ ($\Gamma^*$) for each genotype.

Values obtained from three such graphs for three individual plants of each genotype was used to calculate the mean values of $R_d$ and $\Gamma^*$.

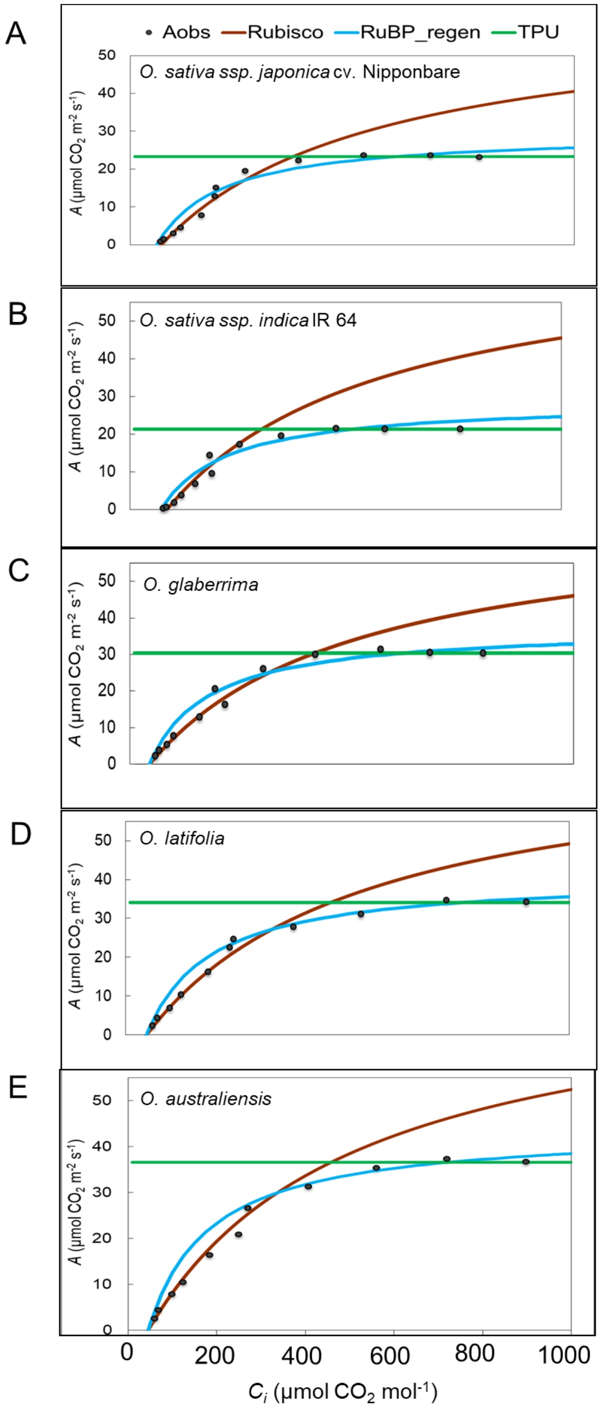


**Fig. S6.** Relationship between $A$ and $C_i$ with $A/C_i$ curve fitting for the selected cultivated and wild rice accessions.

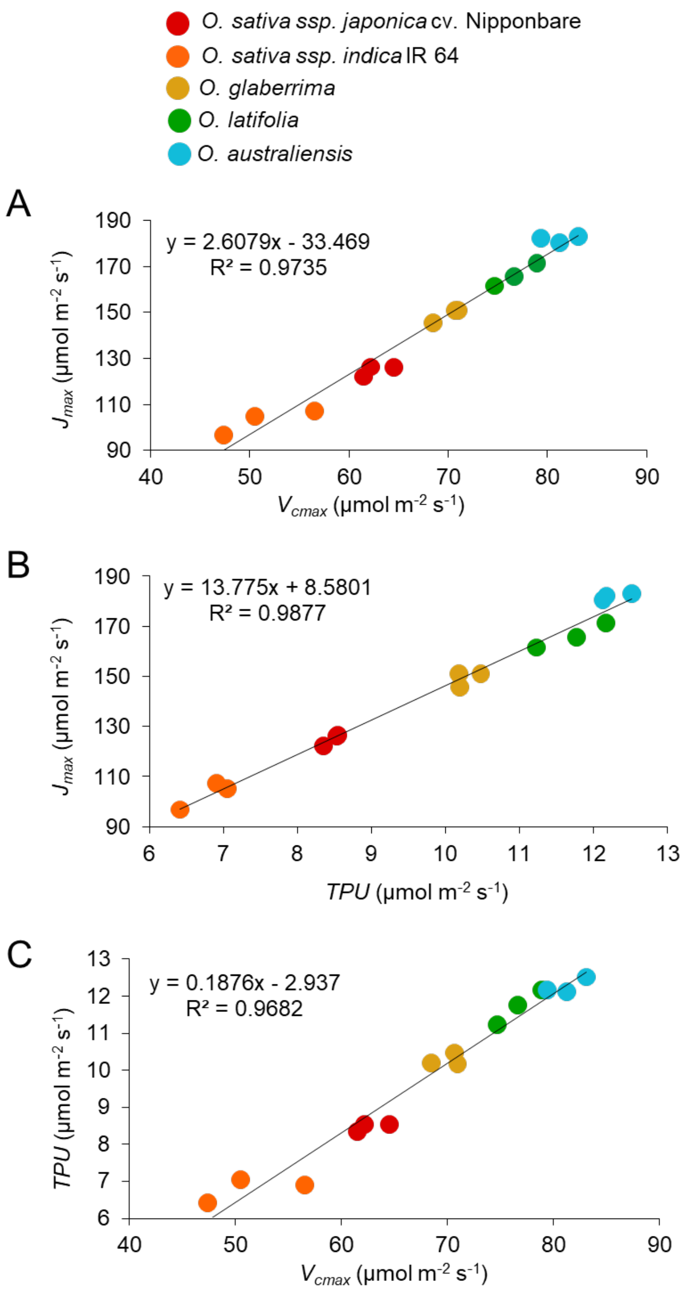


**Fig. S7.** Linear regression analysis between $V_{cmax}$ and $J_{max}$ (A), *TPU* and $J_{max}$ (B), and $V_{cmax}$ and *TPU* (C).

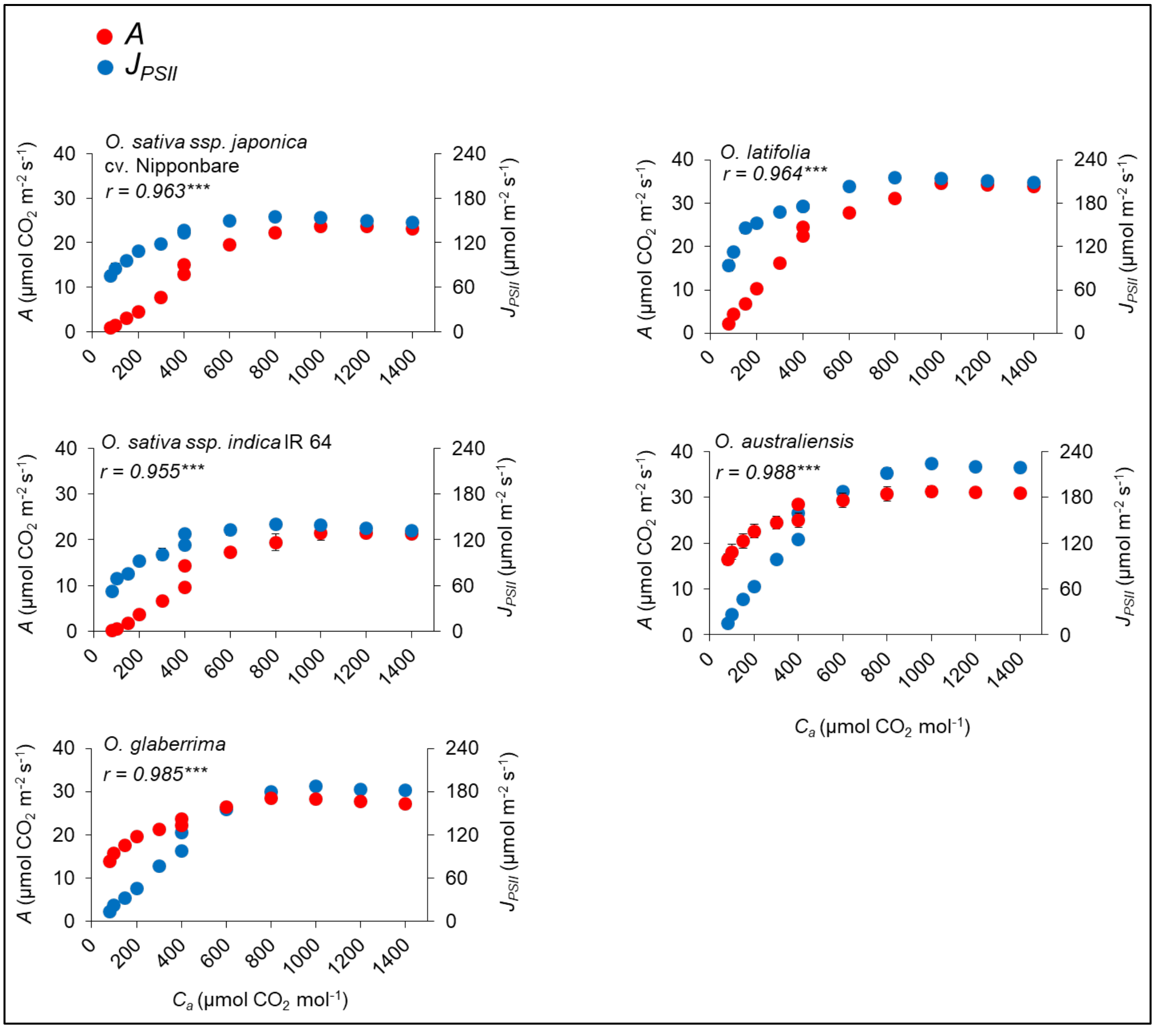


**Fig. S8.** Response and correlation of leaf photosynthesis (*A*) and electron transport rate ($J_{PSII}$) at varying $CO_2$ concentrations for the selected rice accessions. *A* and $J_{PSII}$ are represented with red and blue colors, respectively. Each value represents mean ± SE, where n = five data points from different plants.

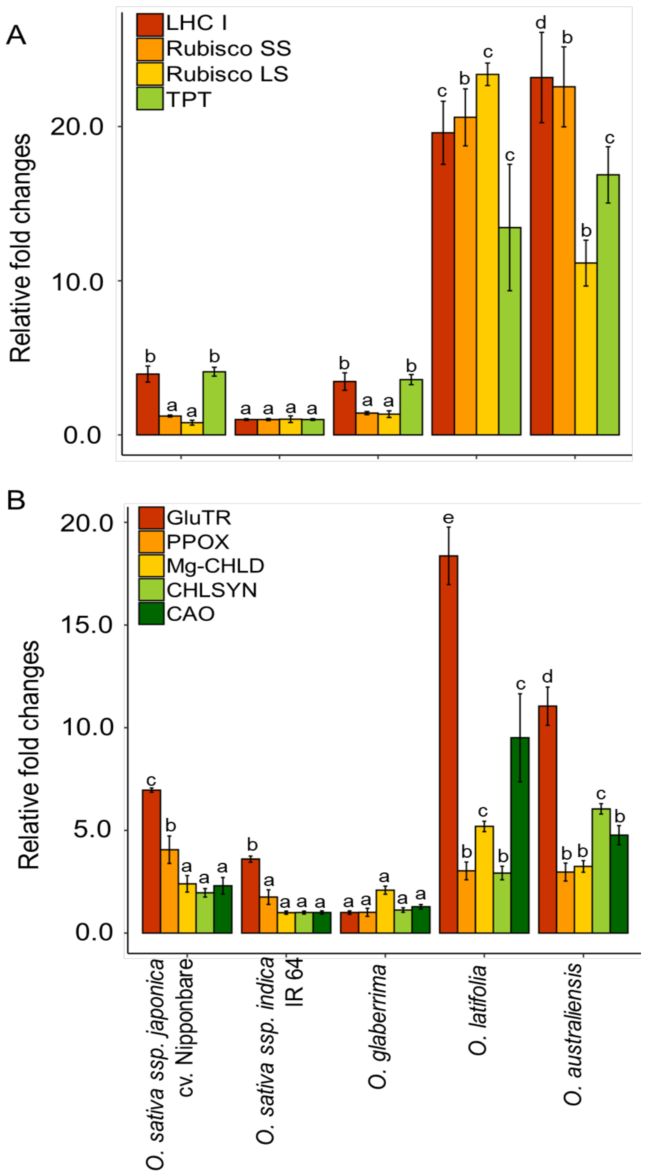


**Fig. S9.** Expression pattern of photosynthetic (A) and chlorophyll biosynthetic genes (B) across the selected accessions normalized to rice ubiquitin (LOC_Os03g03920) internal control.

Values represent mean ± SE (three data points from different plants). Different letters indicate statistical significance according to one-way ANOVA followed by post-hoc Tukey HSD calculation at $P \leq 0.05$.

**Gene abbreviations:** LHCI, Light-harvesting chlorophyll protein complex I; Rubisco SS, Ribulose bisphosphate carboxylase small subunit; Rubisco LS, Ribulose bisphosphate carboxylase large subunit; TPT, Triose phosphate/phosphate translocator; GluTR, Glutamyl-tRNA reductase; PPOX, Protoporphyrinogen oxidase; Mg-CHLD subunit ChlD, Magnesium chelatase; CHLSYN, Chlorophyll synthase; CAO, Chlorophyllide a oxygenase.

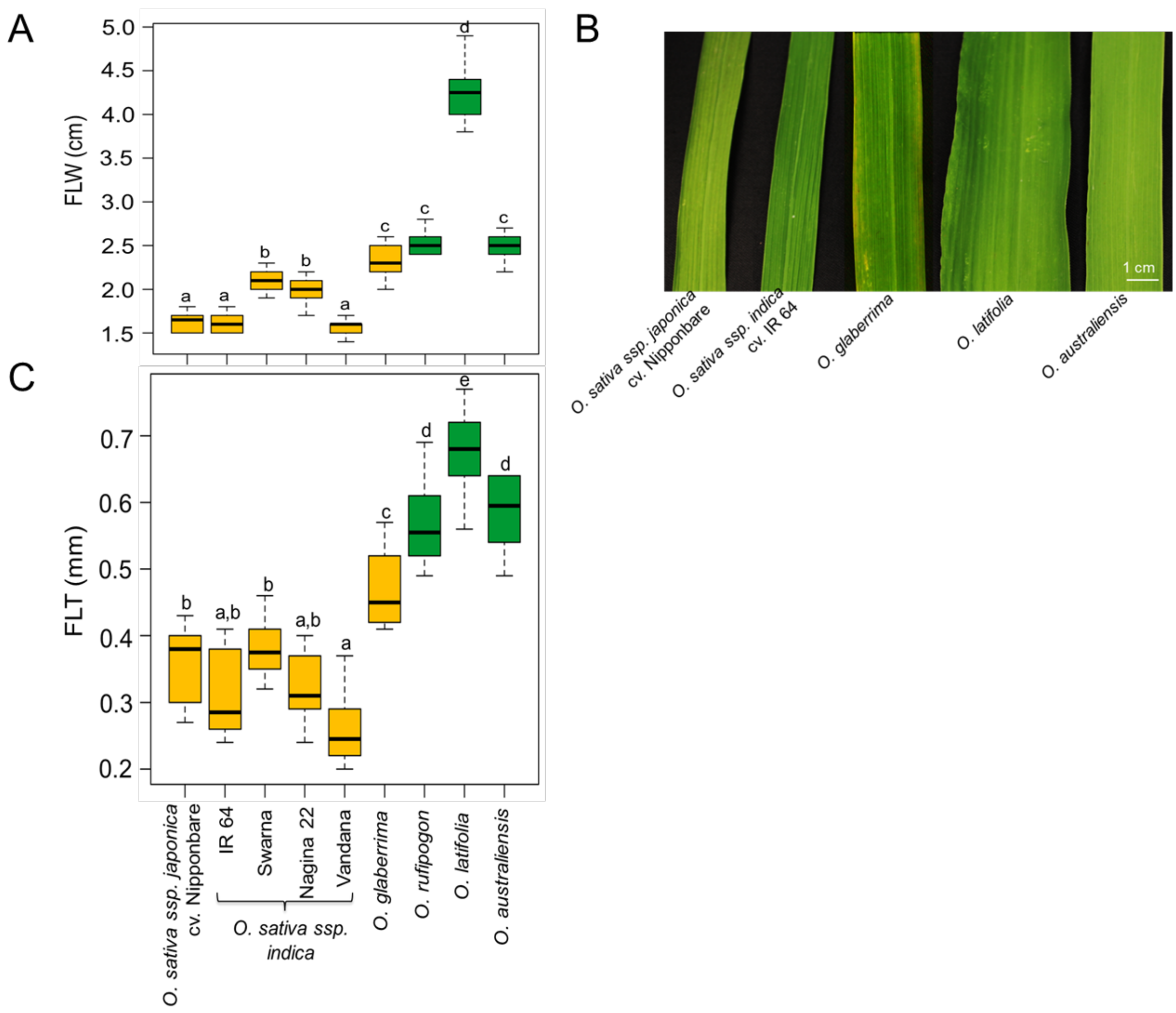


**Fig. S10.** Flag leaf width (FLW, A) and flag leaf thickness (FLT, B) of six cultivated rice varieties and three wild rice species. Also shown are representative images of flag leaves of the five selected cultivated and wild rice accessions (C). Cultivated rice accessions and wild species are represented with yellow- and green- shaded box plots, respectively. Each box plot shows the median and interquartile range. Different letters indicate statistical significance according to one-way ANOVA followed by post-hoc Tukey HSD calculation at $P \leq 0.05$.

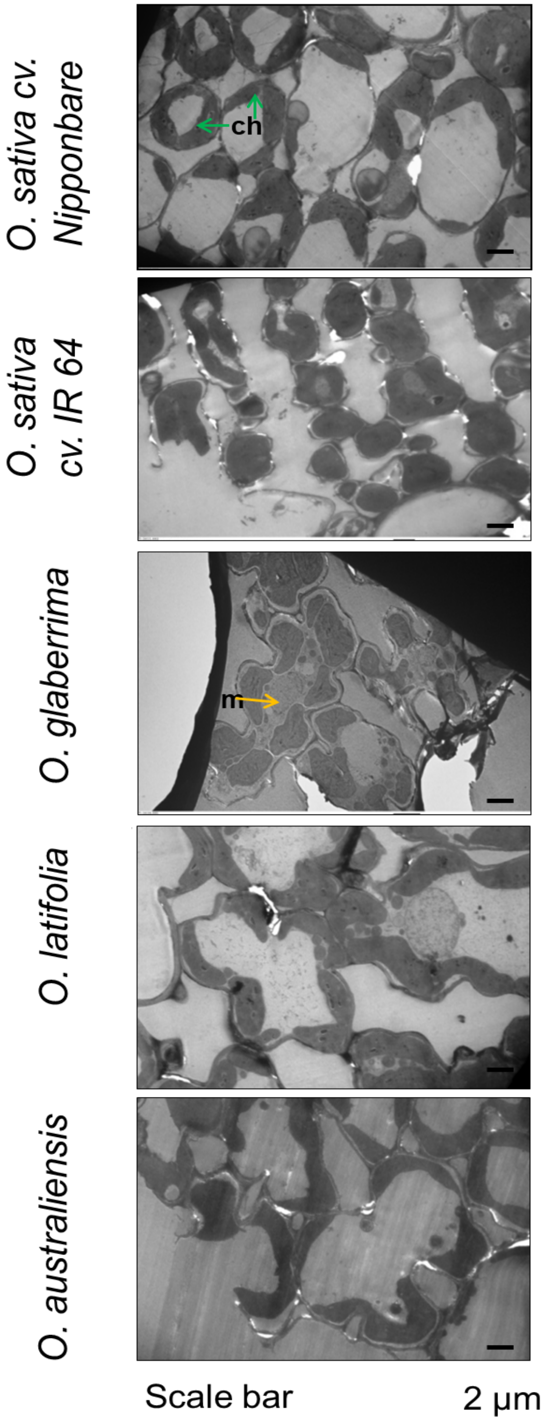


**Fig. S11.** TEM images for the organization of chloroplasts in mesophyll cells. Arrows indicate chloroplast (ch) and mitochondria (m). Scale bar represents 2.00 μm at 1000X magnification.

**Table S1.** Details of rice accessions used in the present study.

| Genome | Genotype | Cultivated /Wild | Cultivar/ accession number | Days to 50% heading |
|---|---|---|---|---|
| AA | *Oryza sativa ssp. japonica* | Cultivated | Nipponbare | 87 |
| AA | *Oryza sativa ssp. indica* | Cultivated | IR 64 | 80 |
| AA | *Oryza sativa ssp. indica* | Cultivated | Swarna | 122 |
| AA | *Oryza sativa ssp. indica* | Cultivated | Nagina 22 | 85 |
| AA | *Oryza sativa ssp. indica* | Cultivated | Vandana | 87 |
| AA | *Oryza glaberrima* | Cultivated | IR 102925 | 89 |
| AA | *Oryza rufipogon* | Wild | IRGC 99562 | 122 |
| CCDD | *Oryza latifolia* | Wild | IRGC 99596 | 89 |
| EE | *Oryza australiensis* | Wild | IRGC 105272 | 101 |

**Table S2.** Total chlorophyll content (*TCC*, µmol $m^{-2}$) and derived leaf absorbance (α) of fully-grown eighth leaves of plants grown under controlled conditions. Each value represents mean ± SE, where n = five data points from different plants. Different letters indicate statistical significance according to one-way ANOVA followed by post-hoc Tukey HSD calculation at $P \leq 0.05$.

| **Genotypes** | **Cultivated/ Wild** | ***TCC*** | ***α*** |
|---|---|---|---|
| *O. sativa* cv. Nipponbare | Cultivated | 470.73 ± 6.11c | 0.86 ± 0.01b |
| *O. sativa* cv. IR 64 | Cultivated | 407.09 ± 13.51b | 0.84 ± 0.01a |
| *O. sativa* cv. Swarna | Cultivated | 513.46 ± 5.10d | 0.87 ± 0.01b |
| *O. sativa* cv. Nagina 22 | Cultivated | 434.36 ± 7.96b | 0.85 ± 0.01b |
| *O. sativa* cv. Vandana | Cultivated | 378.10 ± 10.54a | 0.83 ± 0.01a |
| *O. glaberrima* (IR 102925) | Cultivated | 506.82 ± 3.35d | 0.87 ± 0.01b |
| *O. rufipogon* (IRGC 99562) | Wild | 534.53 ± 11.20e | 0.88 ± 0.01c |
| *O. latifolia* (IRGC 99596) | Wild | 540.67 ± 18.00e | 0.88 ± 0.01c |
| *O. australiensis* (IRGC 105272) | Wild | 580.34 ± 12.79f | 0.88 ± 0.01c |

**Table S3.** Details of genes used for gene expression analysis.

| Gene | Acronym | Locus ID | Function | [1]Pathway |
|---|---|---|---|---|
| Light-harvesting chlorophyll protein complex | LHC I | LOC_Os07g38960 | Functions as a light receptor | *A* |
| Ribulose bisphosphate carboxylase | [2]Rubisco SS<br>[3]Rubisco LS | LOC_Os12g17600.1<br>OrsajCp033 | Catalyzes the rate-limiting step of CO2 fixation in photosynthesis | |
| Triose phosphate/phosphate translocator | TPT | LOC_Os01g13770.1 | Mediates the export of stromal triose-P into the cytosol in a counter exchange for cytosolic Pi | |
| Glutamyl-tRNA reductase | GluTR | LOC_Os10g35840.1 | Synthesis of 5-Aminolevulinic acid (ALA) | *CB* |
| Protoporphyrinogen oxidase | PPOX | LOC_Os01g18320.1 | Involved in the production of heme molecule | |
| Magnesium chelatase subunit ChlD | Mg-CHLD | LOC_Os03g59640.1 | Insertion of Mg into protoporphyrin IX in an ATP-dependent manner | |
| Chlorophyll synthase | CHLSYN | LOC_Os05g28200.1 | Conversion of chlorophyllide to chlorophyll a | |
| Chlorophyllide a oxygenase | CAO | LOC_Os10g41760.1 | Oxidation of chlorophyll a to chlorophyll b | |

[1]Pathways: *A* – Photosynthesis; *CB* – Chlorophyll biosynthesis
[2]Rubisco SS– Small subunit, (nuclear)
[3]Rubisco LS– Large subunit, (plastidial)

**Table S4.** List of primers for qRT-PCR analysis.

| Acronym | Primer pairs (5'-3') | |
|---|---|---|
| | **Forward** | **Reverse** |
| LHC I | CATCTTTCCCCAACAACAAGTTC | GTGAAGGCCTGGAAAATGGTG |
| Rubisco SS | ACTCCAGCTTCGGCAACGTCAGCA | ATACGGACGAATGCATCAGGGTAC |
| Rubisco LS | ACACTGATATCTTGGCAGCATTCCGAG | GTAGAGCGCGTAGGGCTTTGAAAC |
| TPT | TGGAGAAGTACCCTGCCTTGATCA | AGTGTGGGCAAACGAGACTGCAAC |
| GluTR | AGGGAGCATCTATTCATGTTG | TGCATCCTTGAACATCCTATC |
| PPOX | CTGAAAGTGAGCTGGTAGAAG | CGAGGAACAATCCATCATAAC |
| Mg-CHLD | ATGCAATTAAAACTGCTCTGCTG | AGCATCATATTGAACTTGGTTAG |
| CHLSYN | GCATTATTTGGAACCCTTACTC | GTCAATCGCTCCTACACATATC |
| CAO | CGGTGACCTGAAAGATGATAC | CATTGGATTCTACCCTCACTAAC |
| Actin | GAAGTGCGACGTGGATATTAG | CAGACACTGTACTTCCTTTCAG |
| Ubiquitin | CTCGCCGACTACAACATCCA | TCTTGGGCTTGGTGTACGTCTT |

**Table S5.** $A_c$-$A_j$ and $A_j$-$A_t$ transitions for the selected genotypes. The transition points are expressed as intercellular $CO_2$ concentration (µmol $mol^{-1}$). Each value represents mean ± SE, where n = three data points from different plants. Different letters indicate statistical significance according to one-way ANOVA followed by post-hoc Tukey HSD calculation at $P \leq 0.05$.

| Genotypes | Cultivated/ Wild | Intercellular $CO_2$ concentration (µmol $mol^{-1}$) | |
|---|---|---|---|
| | | $A_c$-$A_j$ transitions | $A_j$-$A_t$ transitions |
| *O. sativa* cv. Nipponbare | Cultivated | 250.0 ± 1.0a | 608.3 ± 8.3b |
| *O. sativa* cv. IR 64 | Cultivated | 240.0 ± 10.0a | 524.0 ± 14.5a |
| *O. glaberrima* (IR 102925) | Cultivated | 306.7 ± 6.7b | 656.7 ± 44.8c |
| *O. latifolia* (IRGC 99596) | Wild | 300.0 ± 1.0b | 713.3 ± 29.6d |
| *O. australiensis* (IRGC 105272) | Wild | 325.7 ± 6.7b | 700.0 ± 17.3c |

**Table S6.** Quantification of leaf developmental traits such as Minor vein width (MiV-W; µm), Minor vein height (MiV-H; µm), total number of veins (TV; count), mesophyll lobes (M-Lo; count), mesophyll length (M-L; µm), mesophyll width (M-W; µm), mesophyll area (M-A; $\mu m^2$), bundle sheath number (BS-No; count), and bundle sheath area (BS-A; $\mu m^2$) of the fully expanded flag leaves of the selected cultivated rice varieties and wild rice species. Each value represents mean ± SE. Different letters indicate statistical significance according to one-way ANOVA followed by post-hoc Tukey HSD calculation at $P \leq 0.05$.

| Genotypes | Cultivated /Wild | Minor vein (Miv) | | Total Veins (TV) | Mesophyll (M) | | | | Bundle sheath (BS) | |
|---|---|---|---|---|---|---|---|---|---|---|
| | | MiV-W | MiV-H | | M-Lo | M-L | M-W | M-A | BS-No | BS-A |
| *O. sativa* cv. Nipponbare | Cultivated | 55.5 ± 1.2b | 69.9 ± 1.1b | 77.3 ± 0.7a | 7.6 ± 0.2a | 22.1 ± 0.7a | 14.1 ± 0.3a | 269.3 ± 12.9a | 22.8 ± 0.2b | 5479.2 ± 127.1c |
| *O. sativa* cv. IR 64 | Cultivated | 48.4 ± 1.6a | 59.7 ± 1.6a | 73.3 ± 0.9a | 7.0 ± 0.2a | 21.7 ± 1.0a | 13.5 ± 0.4a | 222.3 ± 10.0a | 18.9 ± 0.1a | 4840.1 ± 132.1a |
| *O. sativa* cv. Swarna | Cultivated | 62.2 ± 1.2c | 71.5 ± 1.7c | 96.3 ± 1.9b | 8.2 ± 0.2a | 22.9 ± 1.1a | 14.2 ± 0.3a | 240.6 ± 11.8a | 24.3 ± 0.2c | 5145.8 ± 113.8b |
| *O. sativa* cv. Nagina 22 | Cultivated | 49.6 ± 1.4a | 56.7 ± 1.5a | 80.0 ± 1.7a | 7.3 ± 0.3a | 25.4 ± 1.3a | 14.3 ± 1.3a | 288.7 ± 10.5a | 19.3 ± 0.2a | 4950.2 ± 393.6a |
| *O. sativa* cv. Vandana | Cultivated | 42.0 ± 2.6a | 62.3 ± 1.7b | 82.0 ± 1.2a | 7.7 ± 0.2a | 24.0 ± 0.8a | 14.1 ± 0.3a | 260.6 ± 8.9a | 18.3 ± 0.2a | 4774.0 ± 131.0a |
| *O. rufipogon* (IRGC 99562) | Wild | 50.9 ± 1.4a | 69.3 ± 1.4b | 111 ± 3.0c | 7.6 ± 0.3a | 29.9 ± 1.1b | 18.9 ± 0.5b | 510.3 ± 17.1b | 24.3 ± 0.2c | 4745.6 ± 139.6a |
| *O. latifolia* (IRGC 99596) | Wild | 59.7 ± 1.6b | 79.5 ± 1.6c | 163 ± 4.5d | 9.1 ± 0.3b | 44.0 ± 2.3c | 25.9 ± 0.6d | 826.8 ± 55.9c | 19.1 ± 0.2a | 5191.9 ± 132.5b |
| *O. australiensis* (IRGC 105272) | Wild | 62.4 ± 1.5c | 88.3 ± 1.7d | 105.3 ± 3c | 7.9 ± 0.2a | 31.4 ± 1.3b | 20.8 ± 0.5c | 592.0 ± 31.7b | 21.4 ± 0.2b | 4656.9 ± 89.9a |
